# Primary production dynamics during the decline phase of the North Atlantic annual spring bloom

**DOI:** 10.1101/2023.05.18.541304

**Authors:** Meredith G. Meyer, Mark A. Brzezinski, Melanie R. Cohn, Sasha J. Kramer, Nicola Paul, Garrett Sharpe, Alexandria K. Niebergall, Scott Gifford, Nicolas Cassar, Adrian Marchetti

**Affiliations:** Department of Earth, Marine, and Environmental Sciences, University of North Carolina at Chapel Hill, Chapel Hill, NC, USA; Marine Science institute and the Department of Ecology, Evolution and Marine Biology, University of California, Santa Barbara, CA, USA; Monterey Bay Aquarium Research Institute, Moss Landing, CA, USA; Division of Earth and Climate Sciences, Nicholas School of the Environment, Duke University, Durham, NC, USA

## Abstract

The second field campaign of the NASA EXport Processes in the Ocean from RemoTe Sensing (EXPORTS) program was conducted in the late spring of 2021 within the vicinity of the Porcupine Abyssal Plain (49.0°N, 16.5°W) in the North Atlantic Ocean. Observations from EXPORTS support previous characterizations of this system as highly productive and organic matter rich, with the majority of primary production occurring in large cells (≥ 5 µm) such as diatoms that are primarily utilizing nitrate. Rates of total euphotic zone depth-integrated net primary production ranged from 36.4 to 146.6 mmol C m^−2^ d^−1^, with an observational period average f-ratio of 0.74, indicating predominantly new production. Substantial variability in the contribution of small (<5 µm) and large cells occurred over the observation period, coinciding with the end of the annual spring phytoplankton bloom. Physical changes associated with storms appear to have impacted the integrated production rates substantially, enhancing rates by ∼10%. These disturbances altered the balance between contributions of the different phytoplankton size fractions, thus highlighting the important role of mixed layer variability in nutrient entrainment into the upper water column and production dynamics. In diatoms, inputs of silicic acid related to deepening of the mixed layer increased silicic acid uptake rates yet concomitant increases in NPP in large cells was not observed. This campaign serves as the high productivity endmember within the EXPORTS program and as such, elucidates how nutrient concentrations and size class play key roles in both low and high productivity systems, but in differing ways.

## 1. Introduction

Oceanic regions experiencing annual spring blooms exhibit high concentrations of biomass and high rates of primary production that are disproportionally important for the global ocean carbon cycle and export. For example, the North Atlantic has been shown to account for 0.55 – 1.94 Gt C yr^−1^ of global carbon export from the euphotic zone with approximately half of that occurring during the annual spring bloom (Sanders et al., 2014; Siegel et al., 2016). Understanding and quantifying the drivers behind these highly productive systems is critical to developing more accurate biogeochemical models and predicting the implications of anthropogenic global climate change. The NASA EXport Processes in the Ocean from RemoTe Sensing (EXPORTS) program was initiated with the goal of advancing these efforts through remote sensing and advanced technology in the upper ocean (Siegel et al., 2016; Siegel et al., 2021). The second field campaign of the EXPORTS program was conducted in the spring of 2021 in the North Atlantic Ocean, coincident with the decline phase of the annual spring bloom in this region.

The North Atlantic has been the study site for many research programs since 1985 due to its highly productive and fairly predictable annual spring phytoplankton bloom. Blooms here have been shown to reach chlorophyll-*a* standing stocks of 1.5-2.0 µg chlorophyll *a* L^−1^, supporting productive surface and deep ocean communities and serving a key role for global carbon export (Frigstad et al., 2015). Because of its ecological and biogeochemical importance and predictability, the North Atlantic has been the focus of large research programs such as North Atlantic Aerosol and Marine Ecosystem Study (NAAMES) campaign, the Joint Global Ocean Flux Study North Atlantic Bloom Experiment (JGOFS NABE), and the 2008 North Atlantic Bloom Experiment (NABE08), as well as the location of long-term monitoring at the Porcupine Abyssal Plain site (PAP). Much of the research that has occurred in this region has focused on better understanding phytoplankton bloom dynamics, specifically the mechanisms behind the initiation of the annual spring bloom and its biogeochemical impacts (Sverdrup, 1953; Behrenfeld, 2010).

Previous studies have unequivocally shown the North Atlantic annual spring bloom to be one of the most productive and extensive phytoplankton blooms in the global open ocean, spanning >2000 km (Siegel et al., 2002; Behrenfeld et al., 2006; Behrenfeld 2010) and a substantial contributor to global carbon export (Henson et al., 2019; Buesseler et al., 2020). However, the exact magnitude of net primary production in the North Atlantic is variable, ranging from 100-210 mmol C m^−2^ d^−1^ (Hartman et al., 2010; Frigstad et al., 2015) with typically over half of annual net primary production in this region occurring during the spring bloom (Garside and Garside, 1993; Koeve, 2001). Commonly used export efficiency metrics, including the fraction of net primary production exported from the euphotic zone, i.e. the carbon export ratio (Ez-ratio; 0.45) and flux transmission below 100 m (T_100_, approximately 1.22 indicating increasing POC flux below the euphotic zone), also vary annually, but indicate that this region experiences larger, more efficient export events than most other oceanic regions, including the Southern Ocean (Buesseler et al., 2020 and references wherein). Previous studies have characterized the phytoplankton community structure as being predominantly diatom and dinoflagellate dominated during the bloom (Warner and Hays, 1994; Henson et al., 2012; Sundby et al., 2016). Additionally, the potential impact of physical drivers, including frequent eddies and storms, on productivity have been characterized at high resolutions (Kortzinger et al., 2008; Hartman et al., 2010). Despite the wide breadth of work already conducted in the North Atlantic, gaps of knowledge (e.g., estimates of gross primary production, f-ratios, and carbon and nitrogen assimilation rates) remain in space and time. Without further synoptic sampling efforts, a comprehensive understanding of the production-export dynamics in the North Atlantic remains incomplete.

A better understanding of and constraint of the annual spring bloom are critical to developing more accurate global carbon models. The PAP region is the second largest oceanic CO_2_ sink on the planet, with some studies estimating flux into the ocean is increasing (1.19 mmol C m^−2^ d^−1^ yr^−1^ from 2002-2016; Macovei et al., 2020) whereas others estimate it is decreasing (Hartman et al., 2012). Regardless, this critical CO_2_ sink is subject to change in coming years as impacts associated with anthropogenic climate change will continue to affect bloom timing and net primary production (Hartman et al., 2012; Behrenfeld et al., 2006). Currently, the magnitude of the spring bloom is believed to be controlled by nitrogen (nitrate; NO_3_ ^−^) availability, leading to exceptionally high carbon to nitrogen ratios (C:N) and carbon overconsumption (i.e., production calculated by carbon assimilation is higher than production calculated by nitrogen assimilation; Kortzinger et al., 2008; Frigstad et al., 2015). Due to a predominance of silica-utilizing diatoms during the beginning of the bloom, silicic acid limitation also appears to be a controlling factor on diatom production, leading to bloom decline, shifting taxonomy, and substantial export of organic matter out of the upper ocean (Cetinic et al., 2015). Resource limitation clearly plays a critical role in bloom extent, and dissipation. However, past and recent studies have revealed the role of physical factors, primarily the mixed layer depth, in controlling light and bloom initiation (Sverdrup 1953; Behrenfeld 2010; Henson et al., 2012). Understanding the relationship between these physical and chemical parameters and how they relate to the phytoplankton dynamics and primary production of the region is critically important as these dynamics are subject to change on both short- and long-term timescales (Capotondi et al., 2012; Benedetti et al., 2021).

The primary goal of the EXPORTS program is to investigate drivers of carbon export in both low and high productivity systems. The second field campaign of the program occurred in May of 2021 where an anticyclonic eddy in the PAP site region was occupied for 31 days in a quasi-Lagrangian manner, collecting a suite of in-situ measurements for primary production, plankton community composition, biogeochemical controls, and grazing rates in order to assess ocean dynamics as they relate to carbon export. This campaign represents the high productivity endmember analysis of the program and was timed to capture the peak and dissipation of the annual spring phytoplankton bloom. This study employs measurements of phytoplankton biomass, biological rate processes, and physical parameters to assess the role of new and regenerated production in temporal primary production trends and carbon export during the decline phase of the bloom. We measured size fractionated rates of DIC, nitrate, ammonium, and silicic acid uptake, and plankton biomass (chlorophyll-*a*, particulate carbon, and particulate nitrogen) to evaluate primary production dynamics over the course of the observation period. This size-fractionated approach allows for the assessment of the roles that phytoplankton of different size-classes play in the ecosystem and of the overall significance of size as a master trait variable in ecosystem processes (Peters 1983; Blanchard et al., 2017). These data combined with the suite of physical, chemical, and biological parameters measured during the EXPORTS campaign enabled us to evaluate these primary production dynamics and their influence on carbon export dynamics at levels not commonly achieved in this region.

## 2. Methods

### 2.1 Sampling Strategy

The North Atlantic Ocean EXPORTS field deployment was conducted from May 1^st^ to May 31^st^ 2021, onboard RRS *James Cook* and RRS *Discovery*, with the majority of measurements analyzed here collected aboard RRS *James Cook* (the exception is ^32^Si-*ρ*Si, which was collected aboard RRS *Discovery*). All sampling occurred within the vicinity of the PAP site (49.0°N, 16.5°W; Fig. 1). The *Cook* sampled following a Lagrangian float that was drogued near 100 m for the entire time, while the *Discovery* sampled a wider geographic area in survey mode. Ship-based sampling occurred as a coordinated effort between in situ sampling with numerous autonomous platforms (i.e., gliders, AUVs, floats) as well as a collaboration with the Ocean Twilight Zone (OTZ) Project on a third ship, B/O *Sarmiento de Gamboa*. Sampling began on May 6, 2021 (Julian Day 126) and ended on May 29, 2021 (Julian Day 149). Autonomous assets were deployed in early April and recovered in late May, extending the period of observation. Due to numerous weather days, a total of 11 sampling days were conducted over 23 days with varying frequency. On days where primary productivity measurements were collected, casts were performed prior to, or coinciding with dawn (cast times ranged from 02:30 to 06:00 UTC) using a standard CTD rosette with 24 20-L spigot Niskin bottles. Seawater was collected at five depths corresponding to the 65%, 20%, 10%, 5%, and 1% of incident irradiance (I_o_). One exception to this sampling scheme occurred on JD 139 (May 19) when only four light depths were sampled, corresponding to 65%, 20%, 10%, and 1% I_o_. The CTD rosette lacked a photosynthetic active radiation (PAR) sensor, so estimates of PAR were derived from midday casts of the Hydroscat’s PAR sensor from the previous day. Euphotic zone depths were defined as the 1% I_o_ depth. Mixed layer depths were defined according to the 0.03 kg m^−3^ density differential measured per CTD cast. All data is publicly available at the SeaWiFS Bio-optical Archive and Storage System (SeaBASS; https://seabass.gsfc.nasa.gov/) website.

**Figure 1.**
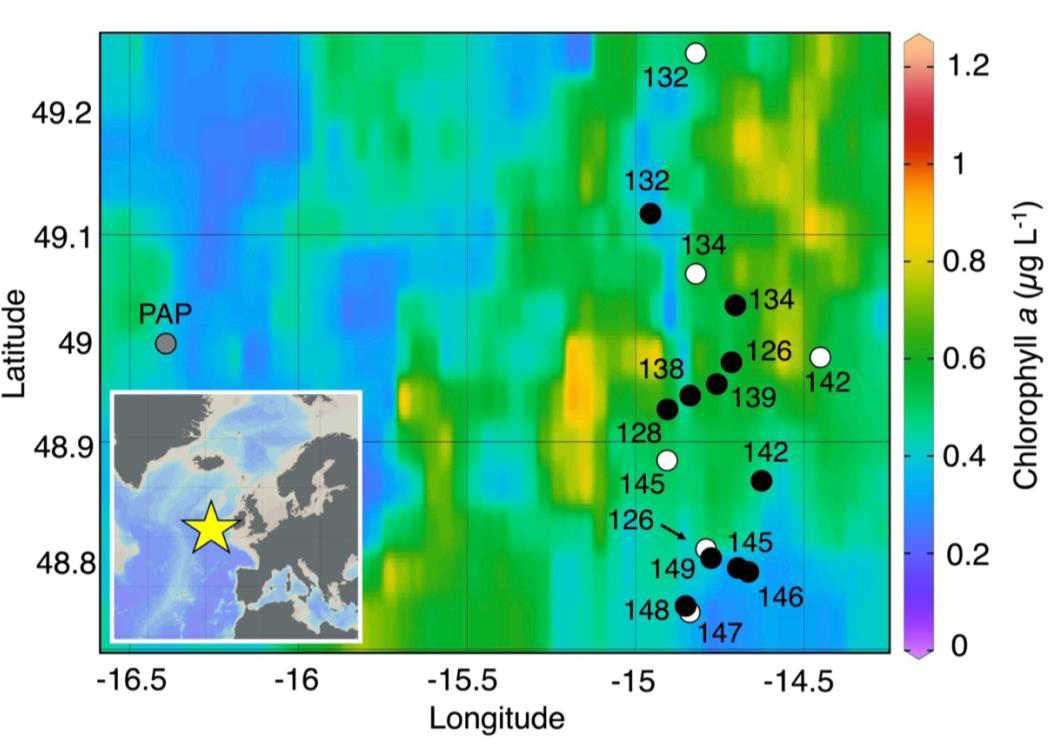
EXPORTS productivity sampling sites in the North Atlantic. Map of sample locations during the North Atlantic EXPORTS field campaign collected aboard RRS James Cook are noted as black dots and identified by Julian day (JD) of sampling, 126-149, corresponding to May 6-29, 2021. Locations of samples collected aboard RRS Discovery are noted as white dots. The Porcupine Abyssal Plain site is noted in gray. Sample sites are overlaid on monthly averaged MODIS chlorophyll a data for May 2021 (https://modis.gsfc.nasa.gov/). The location of the PAP site and our sample sites in relation to the European coast are indicated by the star in the inset map.

### 2.2 Nutrients

Samples were collected for dissolved nutrients (nitrate (NO_3_ ^−^), phosphate (PO_4_ ^3−^), and silicate (Si(OH)_4_)). Dissolved nutrient samples were collected throughout the euphotic zone. Nutrients were analyzed by a Seal Analytical continuous-flow AutoAnalyzer 3 at the UC San Diego Scripps Institute of Oceanography Oceanographic Data Facility according to Gordon et al. (1992), Hager et al. (1972), and Atlas et al. (1971). Ammonium (NH_4_ ^+^) samples were analysed on ship according to the orthophthaldialdehyde (OPA) fluorescence method (Holmes et al., 1999; Taylor et al., 2007).

### 2.3 HPLC Pigments and Size-Fractionated Fluorometric Chlorophyll *a*

Two-liter whole seawater samples were collected and filtered on to pre-combusted GF/F filters for High Performance Liquid Chromatography (HPLC) analysis of phytoplankton pigments. HPLC samples were analyzed according to Van Heukelem and Thomas (2001) and Hooker et al. (2009) at the Ocean Ecology Laboratory at NASA Goddard Space Flight Center (GSFC). See Supplemental Table 1 for the common phytoplankton groups associated with each pigment. A weighted Diagnostic Pigment Analysis (DPA) with the HPLC data was conducted to separate phytoplankton contributions to Chl *a* by size class (micro-, nano-, and picophytoplankton) using weights and ratios determined by Chase et al. (2020) for the North Atlantic. The DPA assumes contributions of different biomarker pigments. Samples for size-fractionated fluorometric chlorophyll-*a* (Chl *a*) concentrations were collected in the same manner as described in Meyer et al. (2022).

### 2.4 Primary productivity and nitrogen uptake rates

Two sets of three 1L seawater samples were collected in acid-cleaned polycarbonate bottles per light depth. One set of bottles was used for size-fractionated, short-term (6 hr) incubations and one set was used for size-fractionated, long-term (24 hr) incubations. Samples for short-term incubations were collected from the upper three depths only with integrations generally confined to the mixed layer whereas samples for long-term incubations were collected from all five light depths. Bottles were acid-soaked (10% HCl) and rinsed with Milli-Q water and seawater prior to and between sampling. Short-term incubation samples were used to estimate gross primary production (GPP), and long-term incubation samples were used to estimate net primary production (NPP) (Marra, 2002; Marra, 2009; Barber and Hiking, 2007).

For primary productivity measurements, bottles were spiked with 334 µmol L^−1^ per 1.07 L (approximately 8% of ambient DIC, assuming DIC concentrations were approximately 1800 µmol L^−1^ and relatively invariant) of NaH^13^CO_3_ isotope. Nitrate uptake measurements employed approximately 10% of ambient Na^15^NO_3_^−^ isotope additions. To calculate additions, ambient NO_3_^−^ concentrations were estimated by an optical nitrate sensor on the profiling Lagrangian float located at the anticyclonic eddy center, and average additions were approximately 16% ambient NO_3_^−^. On four out of eleven sample days, one out of three long-term incubation bottles were spiked with ^15^N-NH_4_Cl isotope to estimate ^15^N-NH_4_^+^ uptake (^15^N-ρNH_4_^+^), approximating 10% ambient NH_4_^+^ concentrations measured in-situ from the previous sample day. All isotope inoculations were added within an AirClean mini flow hood. Following inoculation, bottles were immediately placed in on-deck, flow-through incubators screened with neutral density screening to represent each irradiance light depth.

Following incubation, one of the triplicate bottles was filtered directly onto a pre-combusted (450 °C for 4.5 hours) GF/F filter (Whatman 0.3 µm porosity due to combustion) and represents our total sample. The other two bottles were gravity pre-filtered onto a 5 µm polycarbonate filter (representing our ≥ 5 µm size fraction) and then vacuum filtered onto a pre-combusted GF/F filter (representing our <5 µm size fraction). The two <5 µm and two ≥5 µm samples were averaged to represent small and large size fractions, respectively. Thus, three independent measurements were collected per incubation, per depth. Comparison between the sum of our small and large fractions and the total sample suggests good agreement (r^2^ = 0.93 for ^13^C-ρDIC and r^2^ = 0.70 for ^15^N-ρNO ^−^) with slight decoupling in higher biomass samples. Filters were stored at −20°C until onshore processing at the University of North Carolina before being sent to the UC Davis Stable Isotope Facility for analysis on a mass spectrometer. The atom % ^13^C, atom % ^15^N, particulate carbon (PC) and particulate nitrogen (PN) results per sample, blank, and standard were returned and used to calculate ^13^C-ρDIC, ^15^N-ρNO ^−^, and ^15^N-ρNH ^+^ according to the methods of Slawyk et al. (1997) and Dauchez et al. (1995). Ambient DIC was not measured during the cruise, so concentrations used in the ^13^C-ρDIC calculations were estimated using salinity measured via the CTD rosette and the method of Parsons et al. (1984). ^15^N-ρNO_3_^−^ calculations were corrected for ambient NO_3_ ^−^ concentration at each depth of sampling. For full details on sample collection and processing, see Meyer et al. (2022).

To estimate rates of new production, ^15^N-ρNO ^−^ rates were converted to C units through the 106C:16N ratio (Redfield, 1958), rather than our cruise average C:N of 5.8 (which was comparable to the Redfield ratio albeit slightly low) due to this ratio being non-algal specific. Regenerated production was estimated via two methods. The first method (measured) occurred on days when ^15^N-ρNH ^+^ was measured, and the second method (calculated) occurred on all sample days when regenerated production was calculated according to Equation 1:

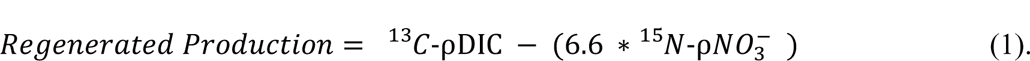

Due to the independent nature of the ^13^C-ρDIC and ^15^N-ρNO ^−^ measurements, it is mathematically possible that 6.6*^15^N-ρNO ^−^ can exceed ^13^C-ρDIC, at which times we assume regenerated production to be negligible. Depth-integrated production rates were calculated via linear extrapolation of the 65% I_o_ sample to the surface and trapezoidal integration of the discrete samples through the deepest sampled depth (1% I_o_), representing the base of the euphotic zone. Uncertainties for the duplicate small and large size-fractions were estimated as the average percent range of the samples and standard deviation, respectively.

A singular observation period-average f-ratio (i.e., the ratio of new to net primary production; Eppley and Peterson, 1979) per size-fraction was calculated via the following equation:

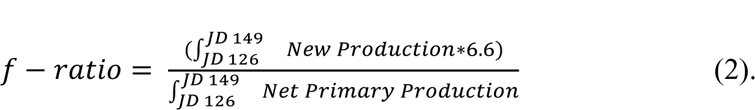

f-ratios were calculated using production rates integrated over the course of the observation period due to the non-steady state nature of the system during time of sampling.

Gross primary production (GPP) rates were calculated by extrapolating the 6 hr ^13^C-ρDIC rates to 24 hours via Equation 3:

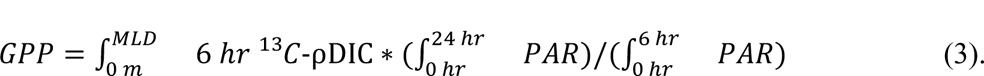

Where 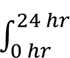 *PAR* is integrated PAR over the 24 incubation and 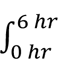 *PAR* is integrated PAR over the 6 hr incubation period as measured via a HOBO PAR Smart Sensor logger attached to the incubator. To account for GPP saturation via photosaturation, levels of saturation (E_k_) were estimated via photosynthesis-irradiance (P/E) curves (J. Fox, personal communication) and the maximum 24 hr PAR was constrained to the maximum PAR that occurred within the 6 hr incubation.

Numerous assumptions are inherent in these productivity calculations. A key assumption is that 6 hr incubations reflect GPP and 24 hr incubations reflect NPP. Previous studies have shown that this assumption does not capture the natural variability in a system (Marra 2002, 2009; Barber and Hiking 2007). Six hr incubations more closely represent a production rate between GPP and NPP, while 24 hr incubations represent a production rate between NPP and net community production (NCP). In Equation 2, we chose to use production rates integrated over the entire sample period because the uncoupling of carbon and nitrate uptake and daily variability made short term f-ratios unrealistic, with values frequently substantially greater than one (maximum f-ratios > 6). Additionally, we chose to use the Redfield ratio to convert nitrogen units to carbon units (Redfield, 1958). Meyer et al. (2022) discuss a more comprehensive list of uncertainties and caveats associated with each of these measurements.

### 2.5 Assimilation rates

Chl *a*-normalized DIC assimilation rates (^13^C-*V*DIC) and Chl *a*-normalized NO_3_ ^−^ assimilation rates (^15^N-*V*NO_3_ ^−^) were calculated by dividing our discrete uptake rates by their corresponding Chl *a* concentrations. Assimilation rates were calculated per size-fraction and total samples and are presented as the average rate within the euphotic zone.

### 2.6 Silicic acid uptake rates

Profiles of silicic acid uptake rates were measured aboard RRS *Discovery* using the radiotracer silicon-32 on six sample days during the observation period (JDs 126, 132, 134, 142, 145, and 147). Seawater was collected from five depths spanning the 55% to the 1% light depths from a trace-metal-clean rosette system. All subsampling of Go-Flo samplers was conducted in a trace-metal-clean van, and the subsamples were transferred within clear plastic bags to a radioisotope van for tracer addition. Seawater for rate measurements was subsampled into trace-metal-cleaned 300-mL polycarbonate bottles and then spiked with 230 Bq of Chelex-cleaned high-specific activity silicon-32 (15,567 Bq µg^•1^ Si). Each sample bottle was capped, and the closure sealed with parafilm before being transferred to deck incubators which were screened for to represent ambient light fields. Incubators were cooled with flowing surface seawater except for those at the two deepest light depths that were held at near in situ temperature using recirculating chillers. Following 24 hr of incubation, water samples were size-fractionated through 25-mm diameter 5.0-µm pore size, and then 0.6-µm pore size, polycarbonate filters. ^32^Si activity on the filters was measured using low-level beta detection as in Krause et al. (2011). Volumetric silicic acid uptake rates (ρ = µmol Si L^−1^ d^−1^) were calculated as in Brzezinski and Philips (1997).

As silicic acid uptake rates were measured on a different ship than other rate measurements, silicic acid uptake rates are matched to data collected aboard RRS *James Cook*, but when silica sample days are offset, environmental parameters (T, S, NO_3_^−^, etc.) measured aboard RRS *Discovery* are included for statistical analyses. Therefore, some temporal and spatial variability still exists between datasets, leading to slight differences in their biogeochemical environments.

Euphotic zone ^32^Si-ρSi was calculated via trapezoidal integration in the same manner as ^13^C-ρDIC and ^15^N-ρNO ^−^. Daily integrated ^32^Si-ρSi was compared to daily integrated ^13^C-ρDIC and ^15^N-ρNO ^−^ to calculate ratios. However, CTDs for ^32^Si-ρSi were offset in deployment time and sample depth from CTDs for ^13^C-ρDIC and ^15^N-ρNO ^−^, thus inherently introducing some spatiotemporal biases. Additionally, ^13^C-ρDIC and ^15^N-ρNO ^−^ from JDs 146 and 148 was averaged to compare to ^32^Si-ρSi on JD 147.

The proportion of NPP attributable to siliceous phytoplankton (most commonly diatoms) was calculated according to Equation 4:

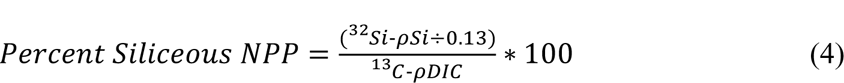

where 0.13 is used to convert mmol Si m^−2^ d^−1^ into mmol C m^−2^ d^−1^ (Brzezinski, 1985).

### 2.7 Statistical analyses of productivity drivers

Z-scores were calculated per day for various environmental parameters (salinity (S), temperature (T), mixed layer depth (Z_MLD_), NO_3_ ^−^, PO ^4−^, Si(OH)_4_, and NH_4_ ^+^), size-fractionated primary production associated parameters (chlorophyll *a*, PC, PN, NPP, new production, ρSi, ^13^C-*V*DIC, and ^15^N-*V*NO_3_ ^−^), and a number of physiologically informative ratios (GPP : NPP, ρDIC : ρNO ^−^, ρDIC : ρSi, ρSi : ρNO ^−^). A heatmap was generated from the z-scores of the daily value (an average per environmental parameter, ^13^C-*V*DIC, and ^15^N-*V*NO ^−^, and ratios and a daily integrated value for Chl *a* and rates) of the above parameters using the “heatmap” function in Matlab 2019a. Dendrograms of these parameters (true values, not Z-scores) were compiled for the EXPORTS PAP site data presented here as well as for data collected during the EXPORTS North Pacific Field Campaign (for details see Meyer et al. (2022) and Brzezinski et al. (2022)). To generate dendrograms, correlation coefficients between cruise wide parameters were first calculated and linkages were calculated based on Euclidean distances between correlations. To reduce potential biases due to lower resolution sampling of ^32^Si-ρSi, we generated two versions of dendrograms-the first dendrograms (A, C) contain all environmental parameters and size-fractionated primary production associated parameters whereas the second dendrograms (B, D) is trimmed to evaluate environmental parameters and size-fractionated ratios from only the days ^32^Si-ρSi was measured. On two sample days in the North Pacific (JDs 234 and 240), ambient NH_4_ ^+^ concentrations were not measured and values were calculated as the average between sample days immediately preceding and immediately following JDs 234 and 240.

## 3. Results

### 3.1 Hydrographic setting

The *Cook* sampled in a quasi-Lagrangian manner in which all sample days occurred within 15 km of the eddy center and with the exception of JD 128, within one water type defined as the eddy surface core water (Johnson et al., preprint). Surface core water is defined as waters located above the subsurface eddy core waters (Johnson et al., preprint). However, it is important to note that within the core water, unresolved lateral variability creates differences in physical (T/S) properties between sample days. Average euphotic zone temperature and salinity were 12.5 °C and 35.55 PSU, respectively. Mixed layer depths (Z_MLD_) measured on the *Cook* were highly variable, ranging from 15 to 197 m over the course of the observation period. Daily peak photosynthetic active radiation (PAR) ranged from 1.75 x 10^5^ to 8.46 x 10^5^ µmol photons m^−2^ s^−1^ over the observation period with an average of 5.94 x 10^5^ µmol photons m^−2^ s^−1^. The depth of the euphotic zone increased from 37 to 59 m from the beginning to end of the observation period, according to measurements collected at noon the day preceding sampling. Four noteworthy storm events (see Johnson et al. for full storm classification) occurred during the observation period on JDs 127-130, 134-135, 138-140, and 141-142 (Johnson et al., preprint). These high wind events led to both the exchange of surface water masses due to horizontal Ekman transports as well as the vertical entrainment of deep waters into the mixed layer from below. Tracer flux analysis suggests entrainment played a larger role on the nutrient budgets of the region than advection did whereas advection had a larger influence on the particulate field (Johnson et al., preprint). Both of these physical processes inherently impacted our results.

### 3.2 Nutrients

Macronutrient concentrations (NO_3_ ^−^, PO_4_ ^3−^, Si(OH)_4_, NH_4_ ^+^) showed consistent temporal patterns within the euphotic zone over the course of the cruise. On average, concentrations were low with slight increases at depth and were punctuated with higher values toward the end of the observation period. Discrete mixed layer NO_3_^−^ concentrations ranged from 4.5 to 6.4 µmol L^−1^ over the observation period with an average (± one standard deviation) of 5.2 ± 0.4 µmol L^−1^. Discrete mixed layer PO_4_ ^3−^ concentrations ranged from 0.3 to 0.5 µmol L^−1^ with an average of 0.4 ± 0.04 µmol L^−1^. Discrete mixed layer Si(OH)_4_ and NH_4_ ^+^ concentrations were nearly depleted at the beginning of the observation period, ranging from below detection limit to 1.6 µmol L^−1^ and 0.1 to 0.8 µmol L^−1^ with average concentrations of 1.0 ± 0.5 µmol L^−1^ and 0.3 ± 0.1 µmol L^−1^, respectively. NO_3_^−^, PO_4_ ^3−^, and Si(OH)_4_ concentrations all increase toward the end of the observation period, following storm 4 (Fig. S1). These increases, which translate to total mixed layer changes of 10-46%, correspond to periods of enhanced wind and mixing of the upper water column which entrained water from below into the mixed layer, increasing nutrient concentrations (Johnson et al., preprint).

### 3.3 Phytoplankton biomass, particulate organic matter, and composition

Consistent with the idea of bloom termination and decline, phytoplankton biomass decreased markedly throughout the euphotic zone by more than 75% over the duration of the observation period. Chlorophyll *a* (Chl *a*) concentrations were initially highest at the start of the observation period, reaching a maximum of 1.8 µg L^−1^ on JD 128 (Fig. 2; Fig. S2). Maximum concentrations were measured at a depth of 36 m and the large size fraction accounted for the majority (86%) of the Chl *a*. Small size fraction Chl *a* was highest (0.46 µg L^−1^) at the beginning of the observation period on JD 126 at a depth of 37 m (Fig. 2A; Fig. S2). However, average total Chl *a* concentrations decreased substantially by 0.40 µg L^−1^ from JD 128-132 following the first storm event of the observation period. Chl *a* concentrations within small and large size fractions were fairly consistent within and below the mixed layer with some elevation (approximately 32 - 64%) at depth at the beginning of the observation period (JD 128) when Chl *a* concentrations peaked (Fig. 2; Fig. S2). These discrete Chl *a* concentrations were slightly lower but within the range of the average Chl *a* concentrations in the eddy center as measured by the Lagrangian float (E. D’Asaro, personal communication).

**Figure 2.**
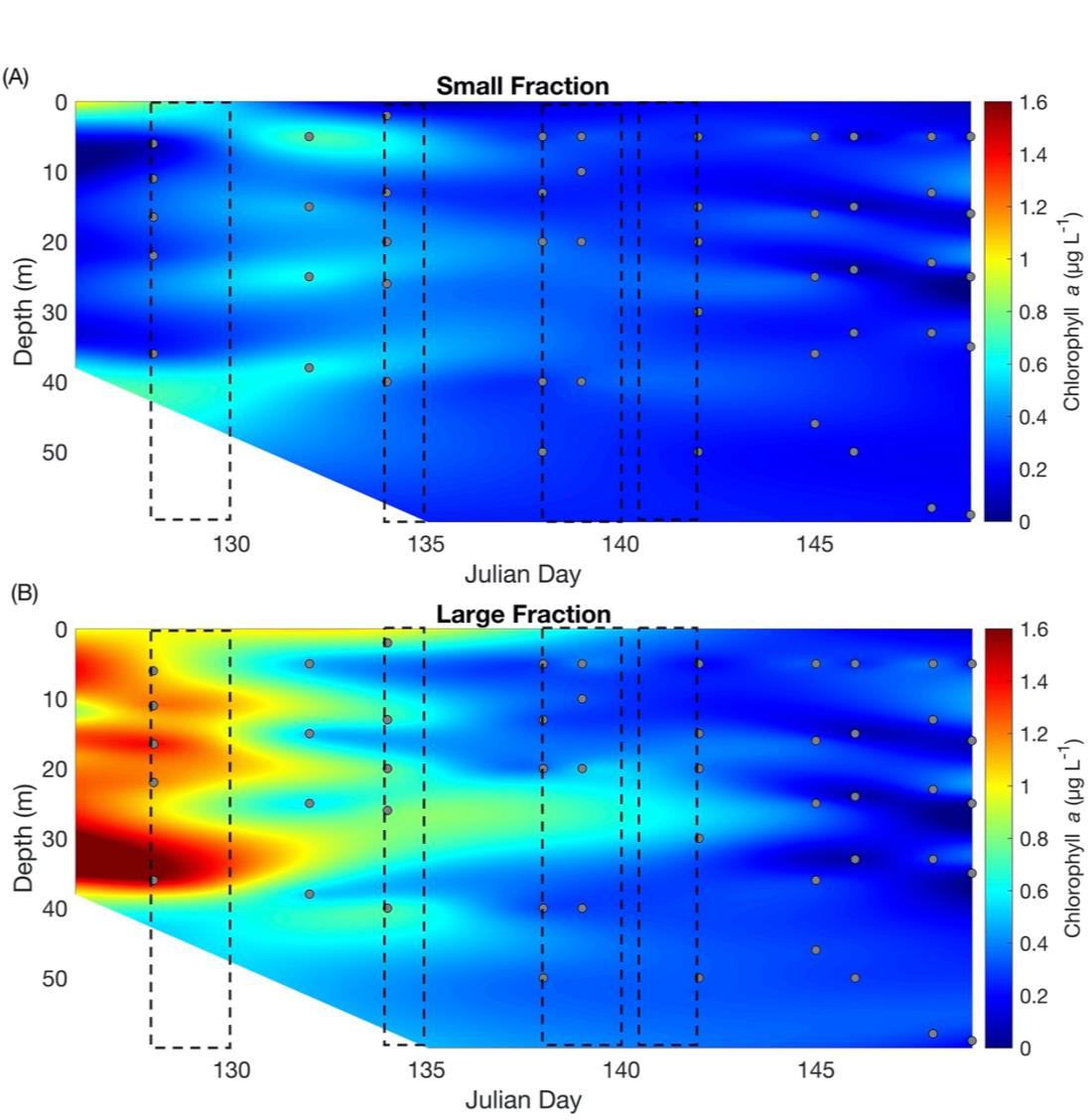
Size fractionated chlorophyll *a* concentrations. Contour plots of the (A) small size-fraction (0.7 - 5 µm) and (B) large size-fraction (≥5 µm) chlorophyll *a* concentrations (µg L^−1^) by depth and Julian day. Grey dots indicate sample depths and white regions indicate an absence of data at those depths. Black boxes indicate storm events. Concentrations between samples are interpolated and thus introduce some uncertainty to those estimates.

Size-fractionated and temporal patterns in particulate carbon (PC) concentrations were similar to those observed in Chl *a* concentration. Large size-fraction PC concentrations were on average higher by 0.17 µmol C L^−1^ with an average concentration of 6.5 ± 4.1 µmol C L^−1^ compared to the small size-fraction average of 6.3 ± 2.1 µmol C L^−1^ (Fig. 3). Temporal patterns between PC and Chl *a* appear fairly consistent, with higher concentrations occurring at the beginning of the observation period before declining around JD 138 and onward. The maximum small size-fraction PC concentration of 13.2 µmol C L^−1^ was observed at 5 m on JD 126, while the maximum large size-fraction PC concentration of 20.6 µmol C L^−1^ deviated from this pattern slightly, occurring at 5 m on JD 148. Total PC ranged from 5.6-27.2 µmol C L^−1^ with an average of 13.7 ± 5.4 µmol C L^−1^. The overall distribution of PC within the water column exhibits slightly divergent patterns compared to those of Chl *a* with higher concentrations in the upper 20 m whereas maximum Chl *a* concentrations occurred at depth, supporting the notion of a subsurface chlorophyll maximum in this region.

**Figure 3.**
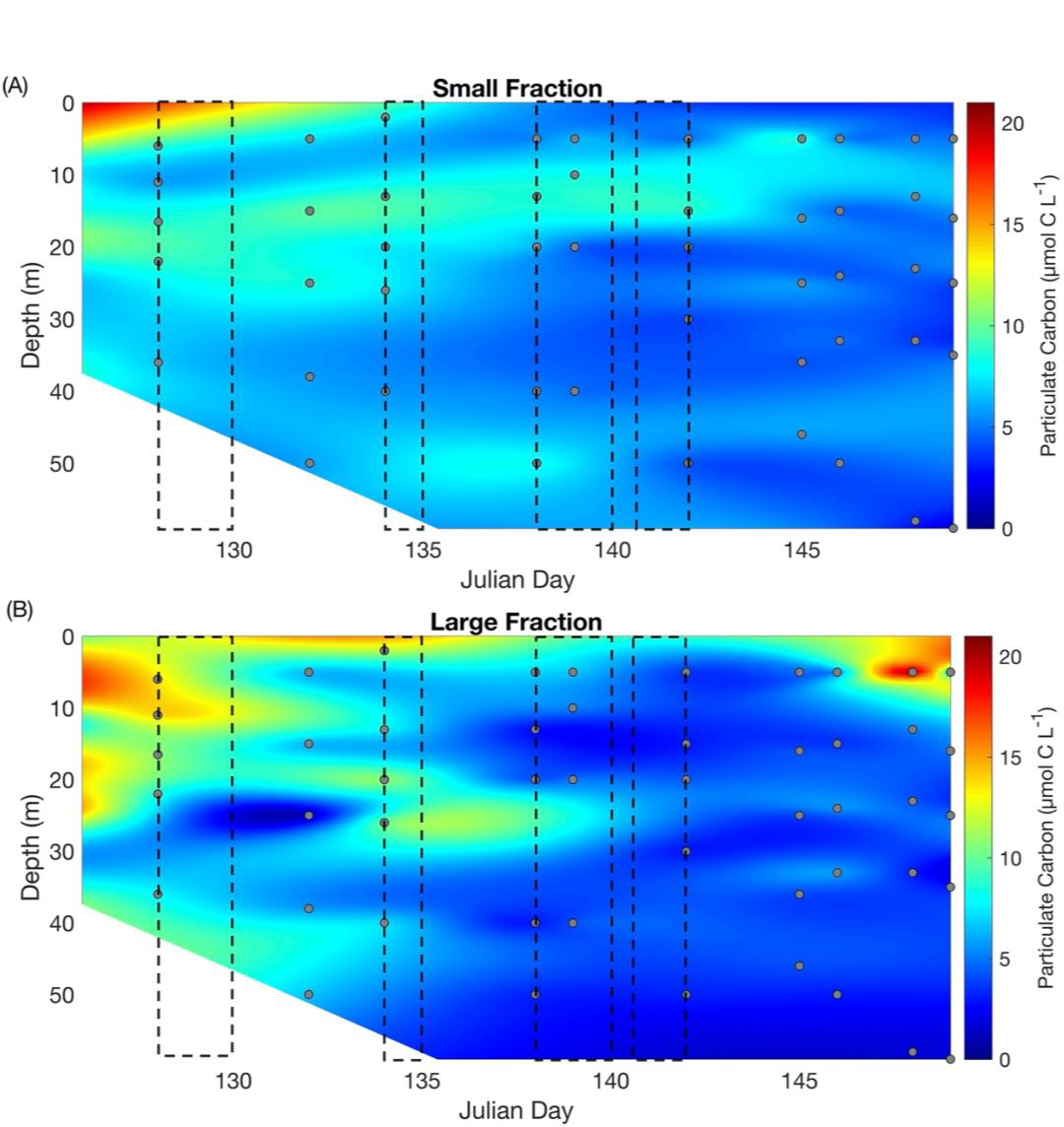
Size fractionated particulate carbon concentrations. Contour plots of (A) small size-fraction (0.7 - 5 µm) and (B) large size-fraction (≥5 µm) particulate carbon (PC) concentrations (µmol C L^−1^) by depth and Julian day. Grey dots indicate sample depths and white regions indicate an absence of data at those depths. Black boxes indicate storm events. Concentrations between samples are interpolated and thus introduce some uncertainty to those estimates.

PC:PN ratios were slightly below the canonical Redfield ratio of 6.6 (Fig. S4; Redfield, 1958). Average ratios were 5.9 ± 1.8, 6.1 ± 3.2, and 5.8 ± 1.2 for the small size-fraction, large size-fraction, and total, respectively. These ratios are low relative to previously reported values (Frigstad et al., 2015), with minimum ratios of 1.1, 1.6, and 2.8 for the small size-fraction, large size-fraction, and total, respectively. Some anomalously high ratios (21.4, 18.6) occurred within the large size-fraction in the upper 15 m over the course of the observation period, but overall, PC:PN ratios were consistent with or lower than the Redfield ratio (122 out of 168 samples were < 6.6).

Total PC:Chl *a* ratios varied drastically over the course of the observation period from 107.7 to 1347 µmol C µg Chl *a*^−1^, with an average of 279.5 ± 237.6 µmol C µg Chl *a*^−1^. The size-fractionated PC:Chl *a* ratios also varied similarly, with values from 101.8 to 1216 µmol C µg Chl *a*^−1^ and 23.6 to 1680 µmol C µg Chl *a*^−1^ for small and large size-fractions, respectively (Fig. S6). Averages were 345.9 ± 250.6 and 257.3 ± 332.2 µmol C µg Chl *a*^−1^ for small and large size-fractions, respectively. Additionally, size-fractionated ratios exhibited temporal trends with extremely elevated ratios in the small size-fraction at the beginning of the observation period (JDs 126 and 128) and extremely elevated ratios in both size-fractions at the end (JDs 146 to 149). In both size-fractions, PC:Chl *a* versus Chl *a* concentrations have a higher correlation coefficient than PC:Chl *a* versus PC concentrations. However, neither PC:Chl *a* versus chlorophyll *a* relationship was statistically significant (R^2^= 0.05 and R^2^ = 0.18 for small and large-fractions, respectively).

HPLC pigment concentrations in the top 55 m show a predominance of fucoxanthin, 19’-butanoyloxyfucoxanthin, and 19’-hexanoyloxyfucoxanthin (Hex-Fuco), accounting for the majority of total primary pigment concentration during the cruise (on average 89%). However, a substantial decrease (approximately 33%) in the relative contribution of fucoxanthin to the summed diagnostic pigments occurs from Julian day 128 to 129, coincident with storm 1 (Fig 4B). During this time, the relative contribution of Hex-Fuco increases substantially, by approximately 30% (Fig 4B). The weighted diagnostic pigment analysis showed a decrease in the microphytoplankton (phytoplankton >20 µm in diameter) over time (Fig 4A). While nanophytoplankton (2-20 µm in diameter) are split between our small and large size fractions using this definition, the microphytoplankton fall within our large size fraction measurements only.

**Figure 4.**
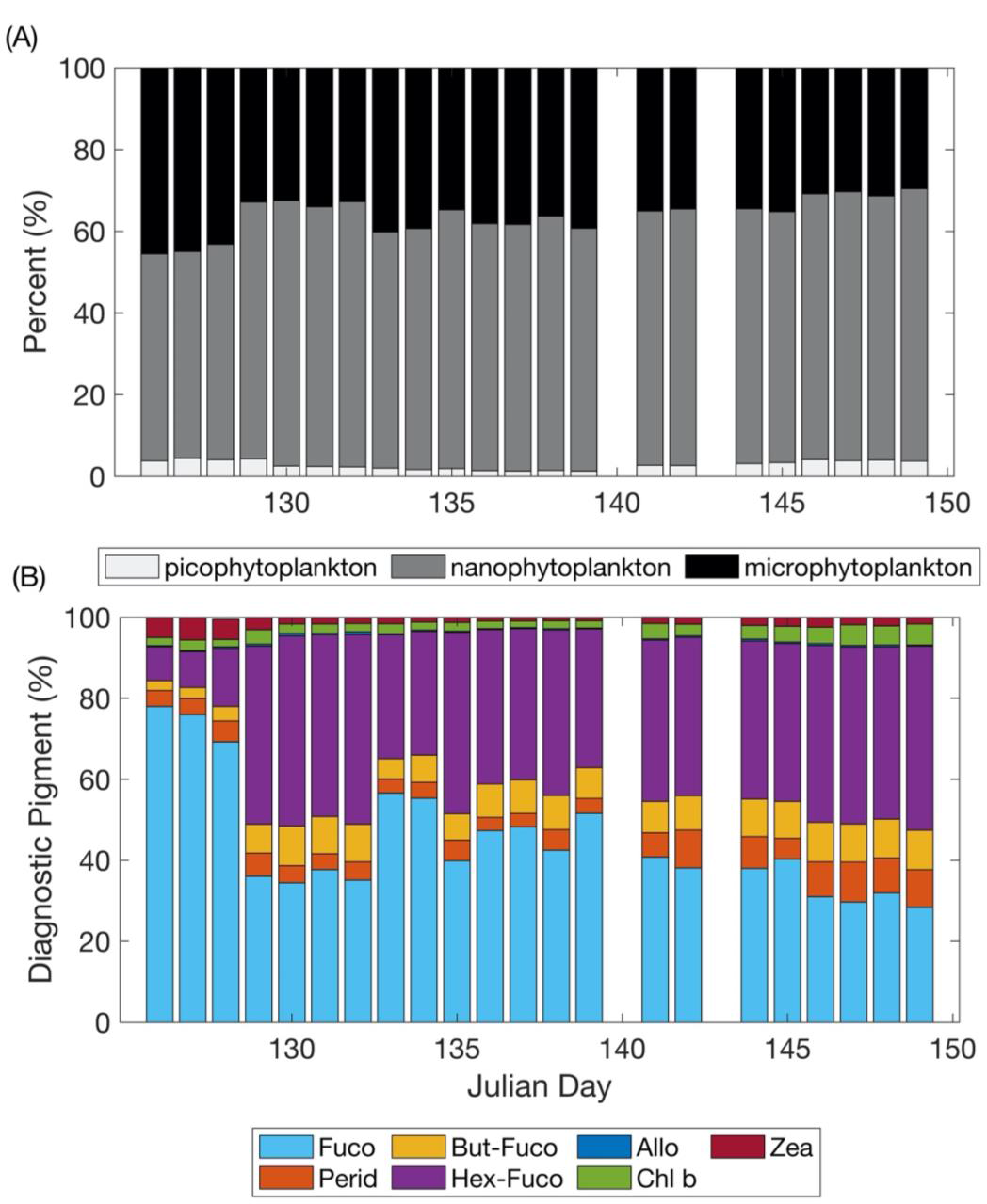
High Performance Liquid Chromatography trends. (A) Fraction of each phytoplankton size class determined by the Diagnostic Pigment Analysis (following Chase et al., 2020) and (B) the percentage (%) of weighted diagnostic pigments to the summed total of all weighted pigments by Julian day. Pigments included in the DPA are: 19’-butanoyloxyfucoxanthin (But-Fuco), 19’-hexanoyloxyfucoxanthin (Hex-Fuco), alloxanthin (Allo), fucoxanthin (Fuco), peridinin (Perid), total chlorophyll b (Chl b), and zeaxanthin (Zea).

### 3.4 Net primary production

Net primary productivity (NPP) displayed temporal trends similar to chlorophyll *a* with a peak in the beginning of the observation period (JDs 126) and a minimum near the end. Size fractionated discrete ^13^C-ρDIC showed higher rates of uptake in the large size-fraction during this time before a switch to higher rates in the small size-fraction following the first storm event on JD 132. From JD 126 to JD 128, the small size-fraction ^13^C-ρDIC was on average 0.81 µmol C L^−1^ d^−1^ and large size-fraction was on average 2.3 µmol C L^−1^ d^−1^ (Fig. 5). On JD 134, the uptake rate for the large size-fraction and total NPP increased throughout the mixed layer with rates comparable to those observed on JD 128 (Fig. 5). From JD 139 to the end of the observation period, uptake for small and large size-fractions were similar, with average values being 0.54 µmol C L^−1^ d^−1^ and 0.28 µmol C L^−1^ d^−1^, respectively (Fig. 5). Total NPP patterns mirrored trends exhibited by the large size-fraction on JDs 126 and 128 before appearing to switch and exhibiting patterns more similar to the small size-fraction from JDs 138 through 149. Total NPP ranged from 0.10-7.3 µmol C L^−1^ d^−1^ with an average of 1.9 ± 1.7 µmol C L^−1^ d^−1^ for the observation period (Fig. 5).

**Figure 5.**
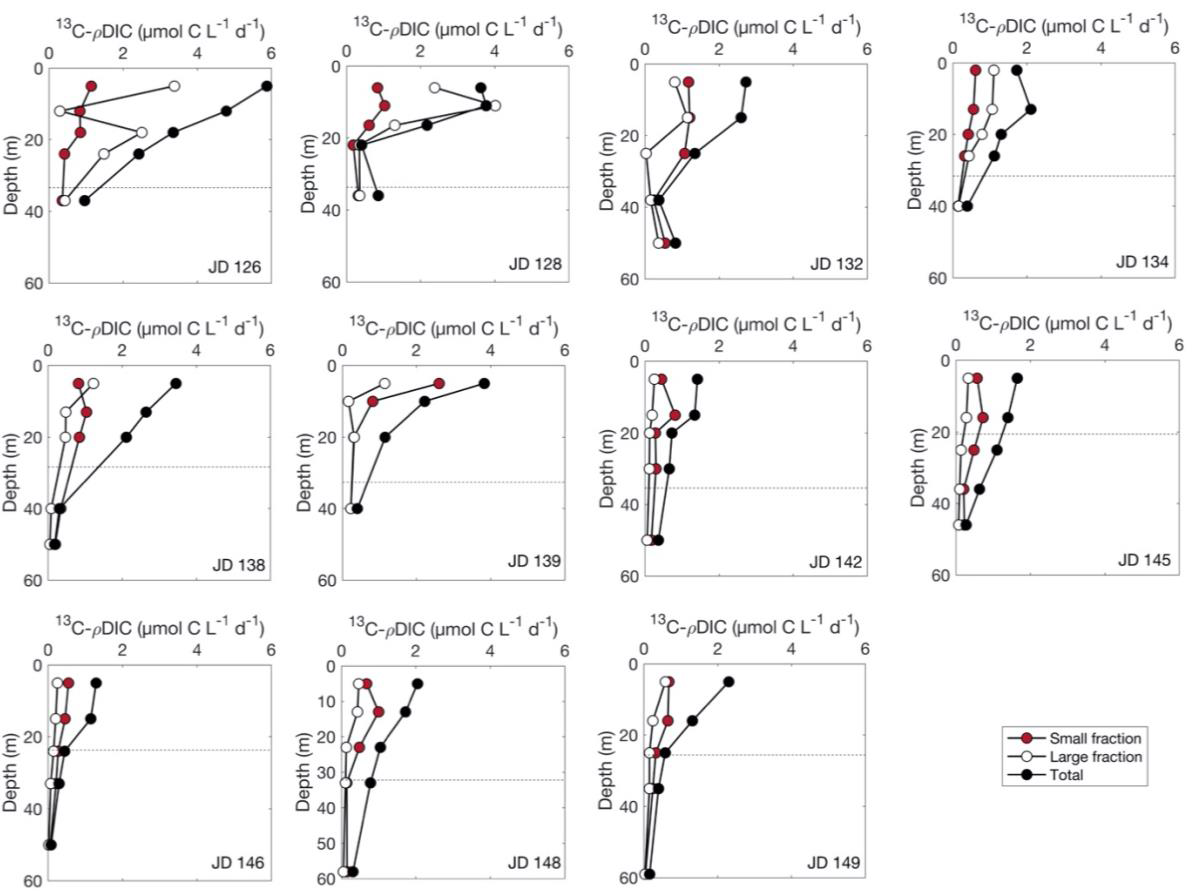
Size fractionated net primary production. Discrete size fractionated carbon-13 isotope uptake (^13^C-ρDIC; µmol C L^−1^ d^−1^) for small (<5 µm), large (≥5 µm), and total particles throughout the euphotic zone by Julian day and depth. Dashed lines indicate mixed layer depths. For the small and large size fractions, the data shown here represents means of duplicate samples. Standard deviations for duplicates were 19.3% and 21.9% of small and large size fractions, respectively.

Total depth-integrated NPP was highest on JD 126 at 146.6 mmol C m^−2^ d^−1^ before declining substantially (Table 1; Fig. 7B). The average total NPP was 82.6 mmol C m^−2^ d^−1^ with minimum values occurring on JD 146 at 36.4 mmol C m^−2^ d^−1^ (Table 1; Fig. 7B). Overall, small and large size-fractions, and total NPP exhibited modest variability and the type of net decline consistent with bloom decline. However, a noteworthy exception to this net decline occurred on JD 134 when NPP reached the second highest rates of the observation period. All size fractions declined over the course of the observation period by 23.7, 74.6, and 88.9 mmol C m^−2^ d^−1^ for small, large, and total NPP, respectively. Total NPP trends appear coupled to trends in the large size-fraction (R^2^ = 0.89) and slightly uncoupled from trends in the small size-fraction (R^2^ = 0.41). A driving factor in the coupling between the large size-fraction and total NPP is the higher concentration of large cells compared to small cells, as evident through size-fractionated Chl *a* concentrations (Fig. 2).

**Table 1.** Daily average chlorophyll *a* (mg m^−2^), particulate carbon (PC; mmol C m^−2^), particulate nitrogen (PN; mmol N m^−2^), net primary production (NPP; mmol C m^−2^ d^−1^), new production (mmol C m^−2^ d^−1^), and regenerated production (mmol C m^−2^ d^−1^) integrated throughout the euphotic zone over the observation period. Dash lines indicate an absence of data on those days. New production is converted to carbon units via the Redfield ratio (6.6C : 1N).

| Julian day | Chlorophyll *a*<br>(mg m<sup>-2</sup>) | Particulate<br>carbon<br>(mmol C<br>m<sup>-2</sup>) | Particulate<br>nitrogen<br>(mmol N<br>m<sup>-2</sup>) | Net<br>primary<br>production<br>(mmol C<br>m<sup>-2</sup> d<sup>-1</sup>) | New<br>production<br>(mmol C<br>m<sup>-2</sup> d<sup>-1</sup>) | Regenerated<br>production<br>(mmol C m<sup>-2</sup><br>d<sup>-1</sup>) | Percent<br>Siliceous<br>NPP (%) |
| --- | --- | --- | --- | --- | --- | --- | --- |
| 126 | 53.1 | 834.3 | 171.5 | 146.6 | 99.4 | 7.9 | 9.0 |
| 128 | 54.1 | 724.2 | 147.6 | 90.5 | 67.1 | -- | -- |
| 132 | 56.7 | 621.5 | 163.0 | 88.0 | 139.5 | -- | 49.8 |
| 134 | 44.5 | 755.2 | 124.0 | 145.2 | 79.9 | 2.9 | 10.6 |
| 138 | 30.3 | 617.4 | 99.8 | 95.4 | 65.5 | -- | -- |
| 139 | 28.8 | 587.0 | 88.3 | 74.8 | 53.0 | -- | -- |
| 142 | 29.8 | 461.2 | 71.5 | 48.4 | 20.2 | -- | 27.9 |
| 145 | 26.9 | 481.1 | 71.6 | 56.6 | 30.6 | -- | 15.6 |
| 146 | 12.7 | 426.2 | 88.2 | 36.4 | 18.5 | 3.5 | -- |
| 147 | -- | -- | -- | -- | -- | -- | 22.3 |
| 148 | 33.1 | 578.7 | 102.0 | 69.5 | 39.4 | -- | -- |
| 149 | 15.5 | 604.4 | 98.5 | 57.7 | 28.1 | 4.4 | -- |
| NA<br>EXPORTS<br>average: | 35.0 | 608.3 | 111.4 | 82.6 | 62.7 | 4.7 | 22.5 |

### 3.5 New production

Daily discrete ^15^N-ρNO_3_ ^−^ was highest in all size-fractions in the surface waters and decreased substantially with depth. Temporal trends between discrete ^13^C-ρDIC and ^15^N-ρNO ^−^ varied slightly with peak total ^13^C-ρDIC occurring on JD 126 at 65% I_o_ whereas peak total ^15^N-ρNO_3_ ^−^ occurred deeper in the water column at 10% I_o_ on JD 128. Most uptake rates showed similar trends between size-fractions with an initial predominance of large cells that shifted over time to a higher relative contribution of small cells later in the observation period.

Changes in discrete ^15^N-ρNO_3_^−^ over the course of the observational period were substantial, reflecting a strong net decline through time. Average small size-fraction rates were 65.9 ± 67.8 nmol N m^−2^ d^−1^, but rates ranged from 2.9-330.3 nmol N m^−2^ d^−1^ (Fig. 6). Average large size-fraction rates over the observational period were slightly higher at 87.8 ± 115.5 nmol N m^−2^ d^−1^ with values ranging from 2.6-534.4 nmol N m^−2^ d^−1^. Average total ^15^N-ρNO_3_^−^ rates were 205.1 ± 208.4 nmol N m^−2^ d^−1^ with values ranging from 5.9-815.0 nmol N m^−2^ d^−1^ (Table 1; Fig. 6).

**Figure 6.**
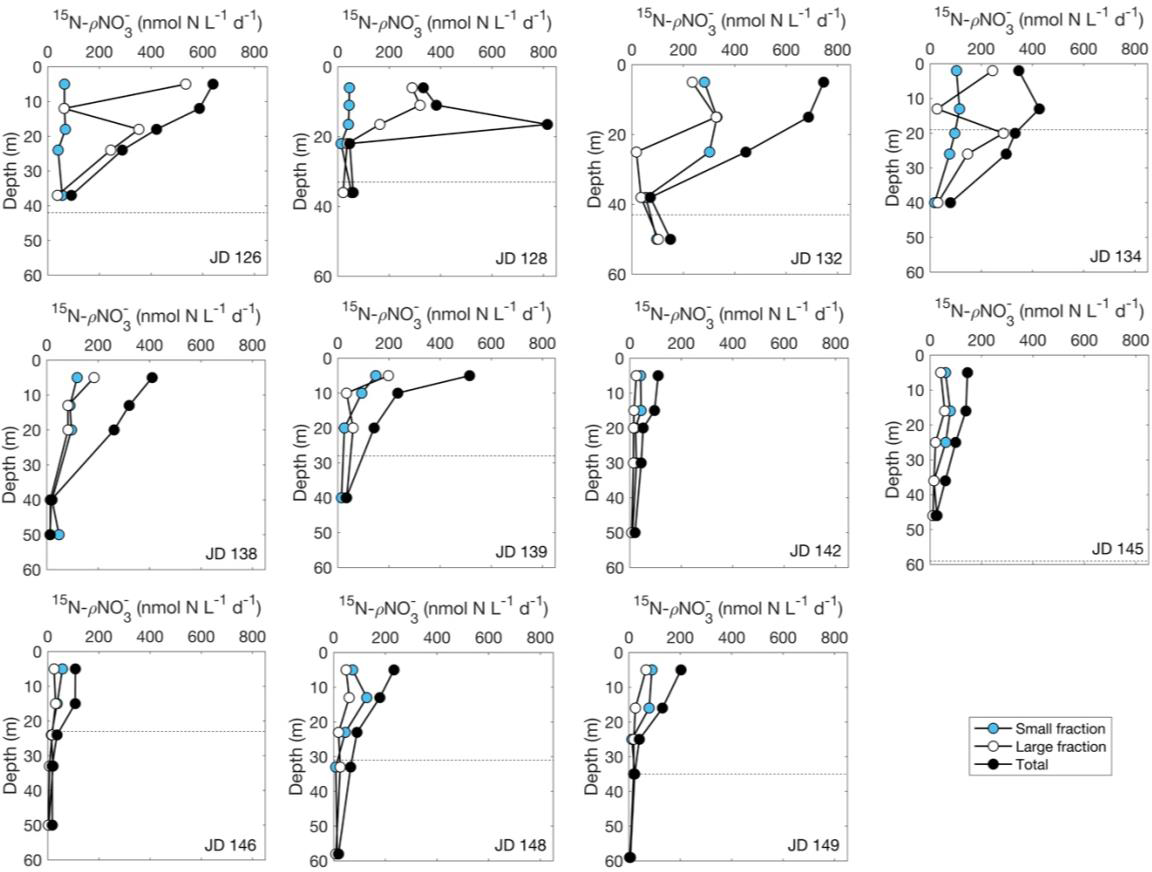
Size fractionated new production. Discrete size fractionated nitrogen-15 (nitrate) isotope uptake (^15^N-ρNO ^−^; nmol N L^−1^ d^−1^) for small (<5 µm), large (≥5 µm), and total particles throughout the euphotic zone by depth and day. Dashed lines indicate mixed layer depths. For the small and large size fractions, the data plotted represents means of duplicate samples. Standard deviations for duplicates were 16.1% and 18.1% of small and large size fractions, respectively.

Magnitudes and trends of size-fractionated rates of depth-integrated new production are correlated (R^2^= 0.63, 0.71) with rates of NPP (Fig 7B,C). Similar to NPP, large size-fraction new production was higher than small size-fraction new production at the beginning of the observation period (JD 126-128) before the trends switched on JD 132, the first sample day since storm 1. Total depth-integrated new production likewise exhibits similar patterns to that of total NPP (R^2^=0.59) with a noteworthy deviation occurring on JD 132. On this sample day, total new production peaked at 139.5 mmol C m^−2^ d^−1^, a rate of 40.2 mmol C m^−2^ d^−1^ higher than the next highest rate of 99.4 mmol C m^−2^ d^−1^ from JD 126 (Fig. 7C). The temporal changes in small size-fraction and large size-fraction new production between peak rates and the end of the observation period exhibited similar magnitudes to those of large size-fraction NPP (56.6 mmol C m^−2^ d^−1^ and 53.4 mmol C m^−2^ d^−1^ for small size-fraction and large size-fraction new production, respectively). However, the decline in total new production from its peak rate on JD 132 to JD 149 was substantially higher than observed with NPP (a decrease of 111.4 mmol C m^−2^ d^−1^).

**Figure 7.**
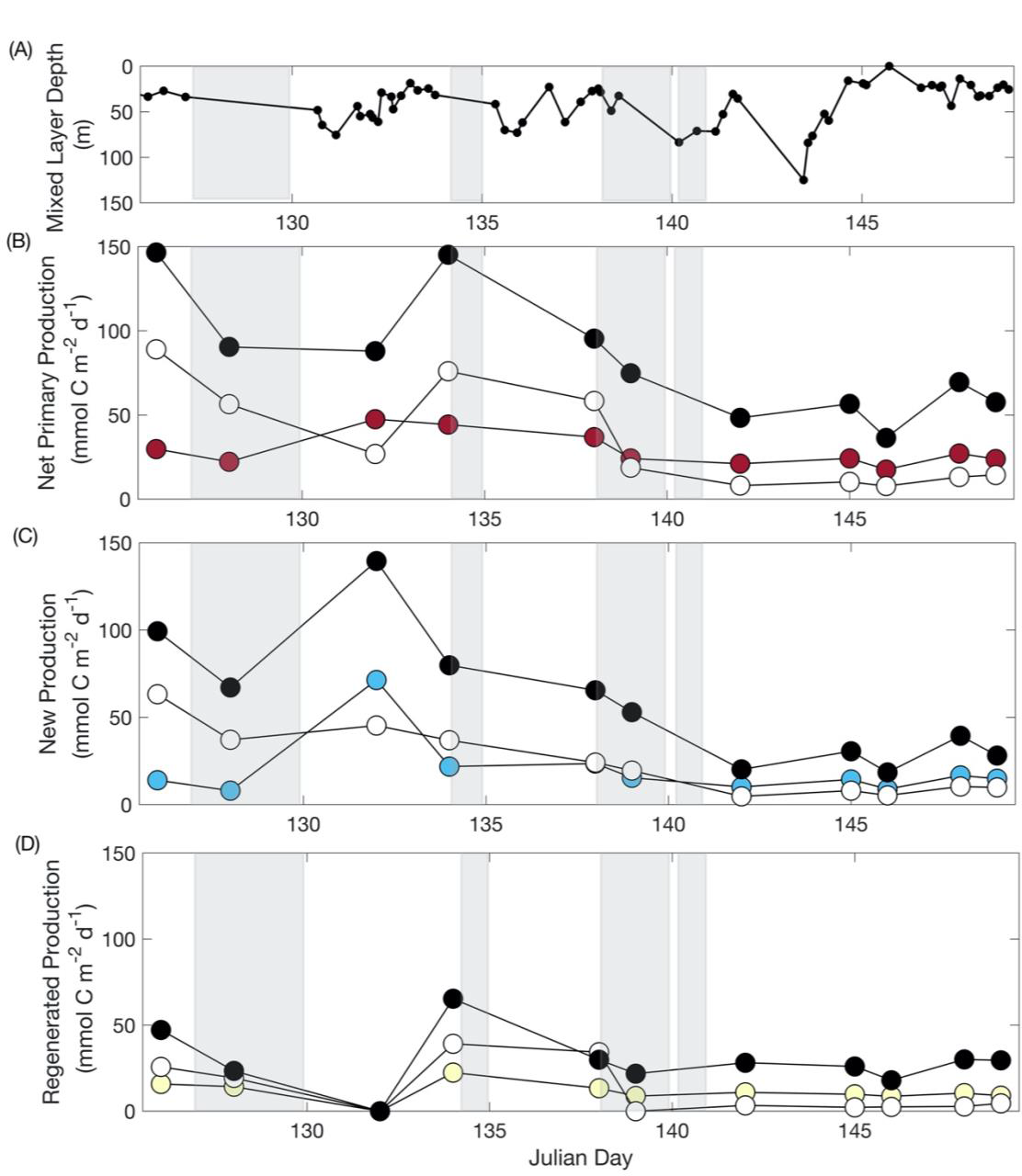
Size fractionated depth-integrated primary productivity. (A) Mixed layer depths and size fractionated (B) net primary production (NPP; mmol C m^−2^ d^−1^), (C) new production (mmol C m^−2^ d^−1^), and (D) regenerated production (mmol C m^−2^ d^−1^) for small (<5 µm), large (≥5 µm), and total particles throughout the euphotic zone by Julian day. New production was converted to carbon units via the Redfield ratio (6.6C : 1N). Shaded regions indicate storm events.

### 3.6 Regenerated production

Regenerated production was calculated according to Equation 1 on every sample day and calculated as ^15^N-ρNH_4_^+^ on four sample days (JD 126, 134, 146, and 149). Average discrete regenerated production rates were comparable between methods, with small and large size-fraction rates of 39.7 nmol N L^−1^ d^−1^ and 44.3 nmol N L^−1^ d^−1^ according to Equation 1, respectively (Fig. 7D). Average regenerated production in the small and large size-fractions from ^15^N-ρNH_4_^+^ were 64.5 nmol N L^−1^ d^−1^ and 43.0 nmol N L^−1^ d^−1^, respectively (Fig. 8). However, the different methods of estimating regenerated production yielded different results in depth-integrated values, with rates calculated from ^15^N-ρNH_4_^+^ on average more than six-times lower than those calculated using Equation 1. The discrepancy in results is likely due to a) calculated regenerated production encompassing more than just ammonium (ie., it encompasses other reduced forms of nitrogen including urea, amino acid, etc. uptake which can be substantial) and b) a decoupling of carbon and nitrogen uptake within incubation experiments. Despite the method used, rates of regenerated production were substantially lower than those of both NPP and new production; total average regenerated production according to Equation 1 was 35.1% of NPP and total average ^15^N-ρNH_4_^+^ was 5.6% of NPP.

**Figure 8.**
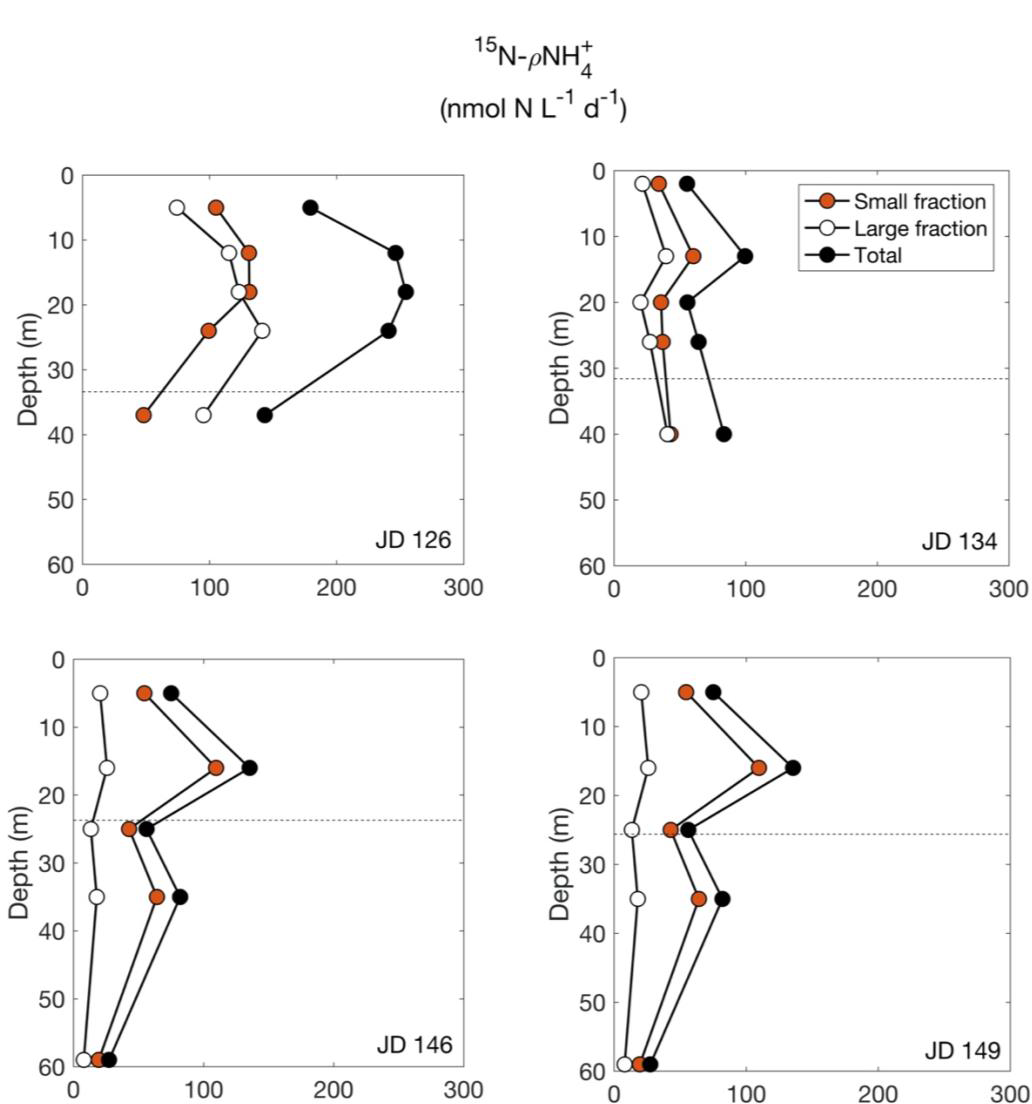
Size fractionated ammonium uptake. Discrete size fractionated nitrogen-15 (ammonium) isotope uptake (^15^N-ρNH ^+^; nmol N L^−1^ d^−1^) by depth throughout the euphotic zone and day. Dashed lines indicate mixed layer depths. Uptake experiments were only conducted on JDs 126, 134, 146, and 149 as represented here.

In contrast to the trends we observed with NPP and new production, regenerated production was comparable between the small size-fraction and the large size-fraction over the observation period and throughout the water column. Regenerated production calculated from Equation 1 had slightly higher rates in the small size fraction whereas rates calculated from ^15^N-ρNH_4_^+^ were slightly higher in the large size fraction. However, the ^15^N-ρNH_4_^+^ method found maximum volumetric rates of regenerated production occurring in the large size-fraction on JD 126 before declining for the rest of the observational period. Small size-fraction regenerated production ranged from 18.9 to 131.3 nmol N L^−1^ d^−1^ whereas large size-fraction regenerated production ranged from 6.8 to 141.5 nmol N L^−1^ d^−1^ (Fig. 8). Total regenerated production ranged from 25.7 to 254.4 N L^−1^ d^−1^, averaging 107.5 ± 70.9 nmol N L^−1^ d^−1^ (Fig. 7). Overall, rates were highest at 10-20 m before declining with depth in through the euphotic zone (Fig. 8).

^15^N-ρNH ^+^ based depth-integrated regenerated production for all size-fractions was highest on JD 126, declined on JDs 134 and 146, and increased slightly on JD 149. Average depth-integrated rates of regenerated production were 2.9, 1.7, and 4.7 mmol C m^−2^ d^−1^ for small and large size-fractions, and total, respectively (Table 1). Integrated small size-fraction regenerated production was higher than that of the large size-fraction on every sample day except at the very start of the observational period on JD 126. Average total depth-integrated NPP was approximately 18-times greater than regenerated production, suggesting regenerated production was only a minor portion of the total production in our study region.

### 3.7 f-ratio

Total average integrated NPP and new production rates were 1950 mmol C m^−2^ d^−1^ and 1450 mmol C m^−2^ d^−1^, respectively over the observation period. Small and large size-fraction rates were comparable at 703 mmol C m^−2^ d^−1^ and 512 mmol C m^−2^ d^−1^compared to 834 mmol C m^−2^ d^−1^ and 579 mmol C m^−2^ d^−1^ for small and large size-fractions NPP and new production, respectively. These values equated to average f-ratios of 0.72, 0.69, and 0.74 for small and large size-fractions, and total, respectively. Average daily f-ratios calculated in the same manner, according to Eq. 2, were comparable but variable. Ratios ranged from 0.36-1.50, 0.41-1.68, and 0.42-1.59 with averages of 0.63, 0.77, and 0.70 for small and large size-fractions, and total, respectively over the observation period.

### 3.8 Gross primary production

Total mixed layer gross primary production (GPP) was high but variable, exhibiting a temporal progression most similar to that of new production with a peak on JD 134, midway through the observation period. The average GPP was 331 ± 198 mmol C m^−2^ d^−1^ with values ranging from 151 to 670 mmol C m^−2^ d^−1^. In contrast to NPP and new production, the majority (51.5% or, on average, 123 mmol C m^−2^ d^−1^) of GPP occurred within the small size-fraction. The small size-fraction exhibited less variable production rates relative to the large size-fraction for 9 out of 11 sample days. Rates ranged from 47.6 to 95.7 mmol C m^−2^ d^−1^. Ratios of GPP : NPP were noticeably higher than the estimated global average of 2.7 (Jones and Halsey, 2015), with averages of 4.5, 3.7, and 4.5 for small, large size-fractions, and total, respectively. These high ratios may suggest a substantial proportion of autotrophic DIC uptake is being lost to respiration over time.

### 3.9 Assimilation rates

Total euphotic zone ^13^C-*V*DIC ranged from 0.2 to 9.5 µmol C µg Chl *a*^−1^ d^−1^ with an average of 2.9 ± 2.5 µmol C µg Chl *a*^−1^ d^−1^ (Fig. 7A). Average small ^13^C-*V*DIC and large size-fraction ^13^C-*V*DIC were 3.0 ± 2.6 µmol C µg Chl *a*^−1^ d^−1^ and 1.6 ± 1.4 µmol C µg Chl *a*^−1^ d^−1^, respectively (Fig. 9A). ^13^C-*V*DIC displayed divergent temporal trends compared to NPP with the highest rates occurring at the end of the observation period on JD 149. ^13^C-*V*DIC in all size-fractions were high at the beginning of the observation period before declining substantially between JD 128 and 132, coinciding with declines in Chl *a* and NPP. Rates increased again on JDs 134-139 but declined thereafter, displaying substantial variability, before a peak on JD 149. Over the course of the observation period, small size-fraction ^13^C-*V*DIC was higher than the large size-fraction ^13^C-*V*DIC (on average by two-fold). At the start of the observation period, small size-fraction ^13^C-*V*DIC was higher than total ^13^C-*V*DIC (0.98-.81 µmol C µg Chl *a*^−1^ d^−1^; Fig. 9A).

**Figure 9.**
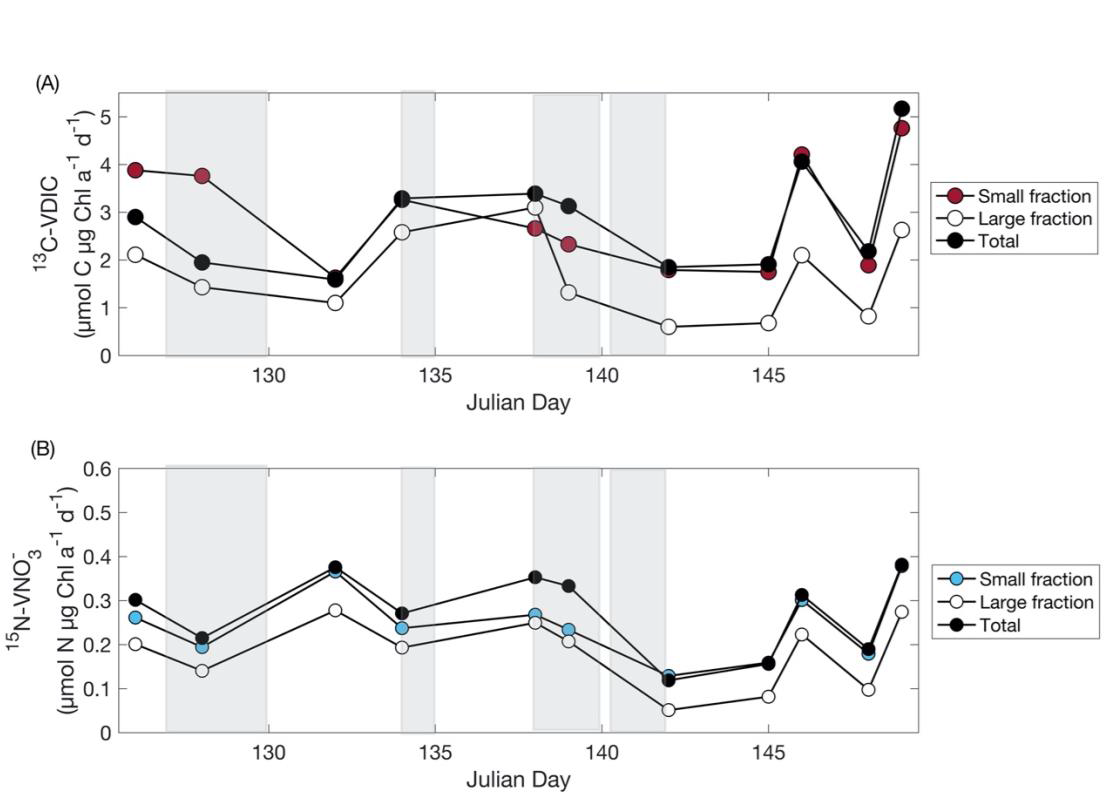
Size fractionated carbon and nitrate assimilation rates. Euphotic zone daily averaged. (A) chlorophyll *a* normalized carbon-13 assimilation rates (^13^C-VDIC; µmol C µg Chl *a*^−1^ d^−1^) and (B) chlorophyll *a* normalized nitrogen-15 assimilation rates (^15^N-VNO_3_ ^−^; µmol N µg Chl *a*^−1^ d^−1^) for small (<5 µm), large (≥5 µm), and total particles by Julian day. Shaded regions indicate storm events.

Total euphotic zone ^15^N-*V*NO_3_ ^−^ ranged from 15.2 – 759.3 nmol N µg Chl *a*^−1^ d^−1^ with average rates of 272.3 ± 221.0 nmol N µg Chl *a*^−1^ d^−1^ (Fig. 7B). Size-fractionated rates were similar, with the average small size-fraction ^15^N-*V*NO_3_ ^−^ of 247.3 ± 188.6 nmol N µg Chl *a*^−1^ d^−1^ and the average large size-fraction ^15^N-*V*NO_3_ ^−^ of 183.1 ± 169.0 nmol N µg Chl *a*^−1^ d^−1^ (Fig. 9B). Toward the beginning of the observation period on JDs 126-128, ^15^N-*V*NO_3_ ^−^ decreased over time, which was a divergence from the pattern in ^13^C-*V*DIC. From JD 134 to the end of the observation period of JD 149, total ^15^N-*V*NO_3_ ^−^ and ^13^C-*V*DIC exhibited similar trends (R^2^ = 0.92) and rates of change (slope = 76.2) as well as similar patterns and rates of change between size-fractions (R^2^ = 0.69 and slope = 66.0 for the small size-fraction, R^2^ = 0.72 and slope = 75.1 for the large-size fraction). Size-fractionated ^15^N-*V*NO_3_^−^ also showed similar patterns to ^13^C-*V*DIC, with higher assimilation rates in the small size-fraction over the observation period.

### 3.10 Silicic acid uptake rates

^32^Si-ρSi ranged from 0.22-4.31 mmol Si m^−2^ d^−1^, 8.81-43.82 mmol Si m^−2^ d^−1^, and 9.77-44.84 mmol Si m^−2^ d^−1^ with averages of 1.99 ± 1.52, 17.75 ± 12.96, and 19.74 ± 12.72 mmol Si m^−2^ d^−1^ for small, large, and total size fractions, respectively (Fig. 10A). The percent siliceous NPP was highly variable in time as well between size fractions. Siliceous phytoplankton consistently accounted for a majority of total NPP in the large size fraction with an average of 93.8 ± 66.6%, but 50% of observations indicate >100% siliceous NPP. These extremely high percentages suggest the 0.13 Si to C conversion factor is unsuitable for this study, likely due to silica limitation within the cells. The relative contributions of small size fraction and total siliceous NPP were both lower, with averages of 6.9 ± 5.3 and 22.5 ± 15.2%, respectively. Given the independent nature of measurements, it is possible that the % siliceous NPP can be large in the size fractionated samples than in the total sample.

**Figure 10.**
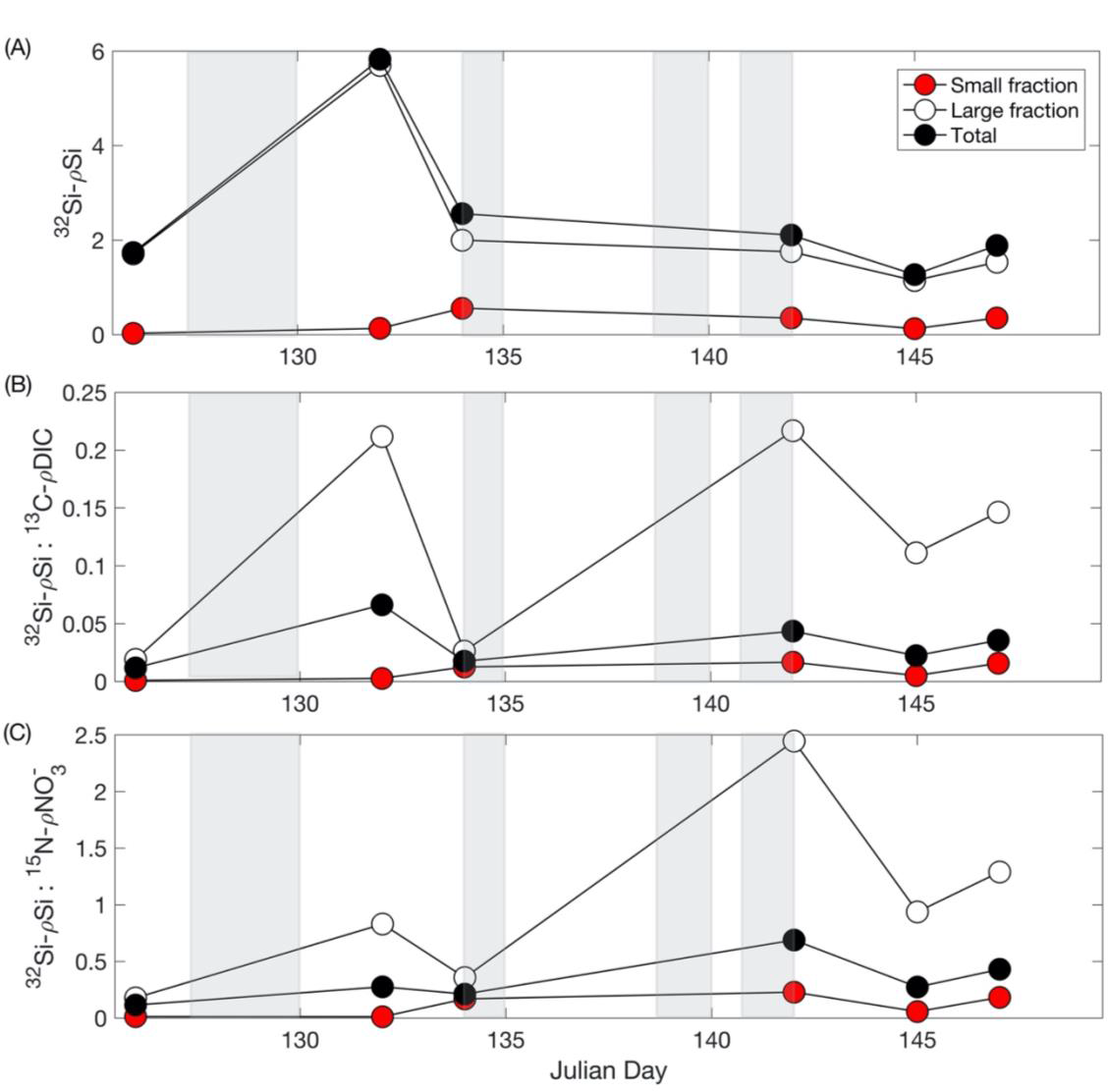
Size fractionated silica uptake ratios and production. Size fractionated ratios of (A) integrated euphotic zone ^32^Si-ρSi, (B) average euphotic zone ^32^Si-ρSi to ^13^C-ρDIC, and (C) average euphotic zone ^32^Si-ρSi to ^15^N-ρNO ^−^ for small (<5 µm), large (≥5 µm), and total particles by day. Shaded regions indicate storm events.

The pattern in total siliceous NPP was fairly consistent with that of new production, with a peak (44.84 mmol C m^−2^ d^−1^) on JD 132 and a decline over time (Table 1). Large size fraction siliceous NPP likewise exhibited a peak (43.82 mmol C m^−2^ d^−1^) on JD 132, while large size-fraction new production did not have a peak on this day. We observe the reverse trend in small size-fraction siliceous NPP, which does not peak until JD 134 (at a value of 4.3 mmol C m^−2^ d^−1^), while new production peaks on JD 132. However, the comparisons between siliceous NPP and new production are limited given the smaller number of calculations of siliceous NPP. Consistently, large size-fraction siliceous NPP accounted for the majority of total siliceous NPP at 88.2 ± 8.6%.

Ratios of ^32^Si-ρSi to ^13^C-ρDIC and ^15^N-ρNO_3_^−^ were calculated and used to assess patterns in silicic acid uptake relative to carbon and nitrogen uptake. The average ρSi : ρDIC was 0.009 ± 0.006, 0.12 ± 0.08, and 0.03 ± 0.02 for small, large, and total size-fractions, respectively (Fig 10). The ρSi : ρNO ^−^ ratios were likewise highly variable with averages of 0.11 ± 0.09, 1.01 ± 0.81, and 0.33 ± 0.20 for small, large, and total size-fractions, respectively (Fig 11B). ρSi : ρDIC and ρSi : ρNO ^−^ showed distinct and similar trends. For both ρSi : ρDIC and ρSi : ρNO ^−^, ratios were highest at the beginning of the cruise (JD 126) before tapering off substantially. While all size-fraction trends were similar, the relationship between ρSi : ρDIC and ρSi : ρNO ^−^ was most similar for the large size-fraction (R^2^ = 0.92, versus R^2^ = 0.62 and R^2^ = 0.80 for small and total size-fractions, respectively). Additionally, the average large size-fraction ratios were much higher (>3x) than the total ratios, while the average small size-fraction ratios were the lowest of all three size-fractions for both ρSi : ρDIC and ρSi : ρNO ^−^, reflecting more siliceous phytoplankton in the large size-fraction than the small size-fraction.

**Figure 11.**
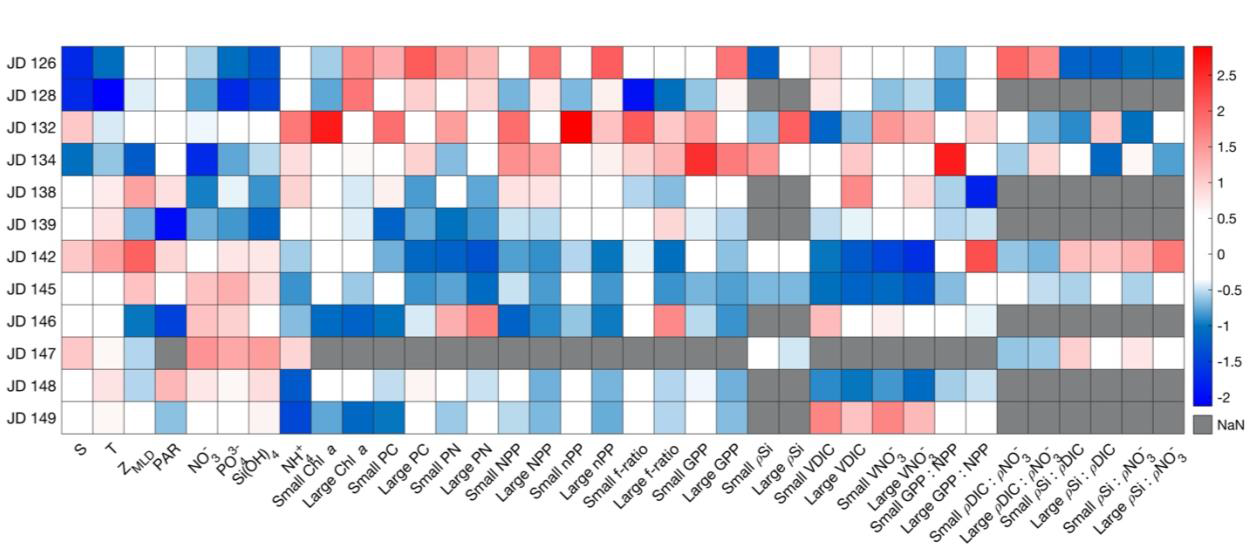
Temporal heatmap of environmental parameters, phytoplankton parameters, and production. Heatmap of Z-scored environmental and phytoplankton biomass, production, and assimilation parameters from the EXPORTS North Atlantic campaign by Julian day.

### 3.11 Statistical analyses of factors influencing primary productivity

The trends in anomalies from the mean values of environmental and production related parameters (Fig. 11) show divergent trends between most environmental parameters and phytoplankton biomass parameters as well as large size-fraction production parameters. At the beginning of the observation period, on JDs 126 and 128, salinity, temperature and macronutrient concentrations were below average as indicated by their negative Z-scores, whereas large size-fraction Chl *a*, small and large size-fraction PC, small and large size-fraction PN, large size-fraction NPP and new production (nPP), and small size-fraction ^13^C-*V*DIC were all either approximately average or above average as indicated by their positive Z-scores (Fig. 11). Silicic acid uptake (ρSi) and the ratio of C to silicic acid uptake (ρDIC : ρSi) were negative, a notable difference from the other production parameters. Interestingly, on JD 132, strong congruent trends in positive Z-scores were observed between Z_MLD_ and small size-fraction chlorophyll *a* and new production. Additionally, small size-fraction PC, NPP, and ^15^N-*V*NO_3_ ^−^ also exhibited positive, albeit not as high, Z-scores. Large size-fraction new production and ^15^N-*V*NO_3_ ^−^ are also moderately positive during this time whereas the other large size-fraction productivity parameters (Chl *a*, PC, PN, NPP, and ^13^C-VDIC) are relatively invariant to negative. Key environmental parameters that would be expected to be associated with enhanced new production, such as PAR, [NO_3_ ^−^], and [Si(OH)_4_] concentrations, are similarly either negative or invariant (Fig. 11).

Another noticeable change in general Z-score trends occurs on JD 142 when all environmental parameters except for NH_4_ ^+^ and ρSi : ρDIC become weakly to moderately positive coinciding with a decrease in all other productivity associated parameters as well as NH_4_ ^+^ (Fig. 11). This change is ephemeral as the Z-scores of some environmental parameters (salinity, temperature, MLD, PAR) decreased substantially on JD 145, and some productivity parameters (small size-fraction PN, ^13^C-*V*DIC and ^15^N-*V*NO_3_^−^, and large size-fraction PN, ^13^C-*V*DIC and ^15^N-*V*NO_3_ ^−^) increase a day later on JD 146. The patterns exhibited from JD 142-146 are consistent with trends of most parameters in the final two sampling days from JD 148-149 (Fig. 11). The divergence between both size-fraction Chl *a* concentrations, PC, PN, NPP, and new production and both size-fraction ^13^C-*V*DIC and ^15^N-*V*NO_3_ ^−^ is noteworthy and has interesting implications for the production dynamics associated with the phytoplankton bloom cessation (see discussion below).

### 3.12 Uncertainty

Coefficients of variation (CV) of the size-fractionated triplicate Chl *a* and duplicate ^13^C-ρDIC and ^15^N-ρNO ^−^ samples was calculated to estimate data variability and uncertainty. Outlier CVs >25% were flagged, assessed, and removed unless high CVs existed across space and time, suggesting the potential that the variability was real. Average Chl *a* CVs were 16.1 ± 14.0% and 18.0 ± 15.7% for small and large size fractions, respectively. Minimum and maximum values were likewise similar across size fractions at 0.00%, 53.6% and 0.8%, 61.9% for the small and large size-fractions, respectively. Across both size fractions of ^13^C-ρDIC and ^15^N-ρNO ^−^, the average CVs were fairly low and consistent at 16.1 ± 12.8% and 18.1 ± 10.4% for small and large ^15^N-ρNO ^−^ and 19.3 ± 17.1% and 21.9 ± 18.1% for small and large ^13^C-ρDIC, respectively. ^13^C-ρDIC had higher maximum CVs of 63.6% and 82.5% for small and large size fractions relative to 49.8% and 43.4% which were observed in the small and large size fraction ^15^N-ρNO ^−^ samples. Minimum CVs were 0.3%, 4.0%, 0.03%, and 0.3% for small size fraction ^15^N-ρNO ^−^, large size fraction ^15^N-ρNO ^−^, small size fraction ^13^C-ρDIC, and large size fraction ^13^C-ρDIC, respectively. The low and consistent average CVs across all datasets lends confidence to data accuracy and reproducibility despite some potential biomass loss associated with the methodology (see below).

While the CV of total ^13^C-ρDIC and ^15^N-ρNO ^−^ samples could not be calculated due to no replicates being collected, the sum of small and large size fraction samples was compared to the total ^13^C-ρDIC and ^15^N-ρNO ^−^ samples to assess biases associated with cell loss. Agreement between ^13^C-ρDIC samples was high with an R^2^ value of 0.89 (Fig. S3A). ^15^N-ρNO ^−^ samples were more variable with a lower R^2^ value of 0.70 and higher values observed within the total ^15^N-ρNO ^−^ relative to the summed ^15^N-ρNO ^−^ (Figure S2B), suggesting losses due to size-fractionation.

## 4. Discussion

### 4.1 Temporal and regional context

The observations presented here demonstrate a system representative of the larger North Atlantic region. Average summertime Chl *a* concentrations from 2003-2015 at PAP were approximately 2 µg L^−1^ (Frigstad et al., 2015) whereas average NPP rates were similarly high at 100-210 mmol C m^−2^ d^−1^ (Hartman et al., 2010; Frigstad et al., 2015). The peak Chl *a* concentrations and NPP rates observed on the EXPORTS cruise were within the range of these previously reported measurements. Average seasonal (February to August) new production is variable, with some studies suggesting that new production ranged from 40.3 to 85.4 g C m^−2^ a^−1^ over a three-year time period (Hartman et al., 2010). Historical estimates of new production are typically calculated according to nitrate budgets (Kortzinger et al., 2008; Hartman et al., 2010) and as such, estimates of short-term rates (as well as those measured via uptake incubations) like those presented here are difficult to compare. Historical estimates of in situ regenerated production from the PAP site are lacking, and thus, this dataset represents a unique comparison of measured and calculated regenerated production estimates from the region.

Data collected as part of the NAAMES and JGOFS NABE campaigns provide useful spatial context for interpreting our EXPORTS findings. The second NAAMES campaign was conducted in late spring 2016, catching the peak and decline of the North Atlantic bloom. Presumably, this event led to enhanced Findings were within the range observed during EXPORTS with average Chl *a* concentrations of 2.3 ± 1.6 µg L^−1^, POC concentrations of 172.6 ± 105.9 mg C m^−3^, and modeled NPP rates of 121.8 ± 36.7 mmol C m^−2^ d^−1^ (Fox et al., 2020). Findings from NABE showed similar rates of average NPP at 90.4 mmol C m^−2^ d^−1^ but suggest a very different balance between new and regenerated production with a lower f-ratio of 0.45 (Martin et al., 1993). This result suggests some degree of spatial continuity in NPP estimates in the larger North Atlantic region (NAAMES took place in the western North Atlantic, while JGOFS NABE was in the eastern North Atlantic but south of the EXPORTS site). While the carbon sequestration potential of the North Atlantic is still debated, our NPP is consistent with the NABE estimates of the 1980s and those of NAAMES in the late 2010s.

The North Atlantic has long been characterized as a region of high primary production and a high export ratio owing to the predominance of large phytoplankton groups such as diatoms, reduced microbial recycling in the surface ocean due to enhanced non-biological removal of carbon, and episodic physical variability (i.e., mixed layer variability) in the region (Henson et al., 2019). Recent work has also quantified a disproportionally large contribution (∼26%) of the mixing pump (i.e., the transport of particulate and dissolved carbon by vertical mixing, eddy subduction, detrainment, and other physical processes; to total export) in the larger North Atlantic region (Nowicki et al., 2022). Carbon sequestration times (i.e., the amount of time before sequestered carbon returns to the surface ocean) for this region are some of the longest reported globally at approximately 128 years (Nowicki et al., 2022). Additionally, the role that new production plays in estimates of total net primary production is high (>70% during the spring season; Garside and Garside, 1992) with increased rates in higher latitude portions of the North Atlantic relative to lower latitude regions (Penta et al., 2021). However, a comprehensive understanding of the relationship between impacts of the substantial mixed layer variability (Nowicki et al., 2022) and new production from phytoplankton size classes as well as within seasonal and episodic time frames is lacking. Our study provides valuable insight into these relationships during the annual spring bloom decline period.

### 4. 2 The impact of a changing physical environment on bloom dynamics

The timing of the EXPORTS field campaign was intended to capture the assumed continuous dissipation of the North Atlantic annual spring bloom. However, the turbulent environment appears to have disrupted what would have been a linear dissipation with nutrient injections and periods of higher sustained assimilation and production than were anticipated. The North Atlantic is known to be a particularly turbulent, stormy system with a high concentration of cyclonic and anticyclonic eddies (Koeve et al., 2002; Waniek 2003; Penta et al., 2021). Given the sustained, high-resolution sampling that occurred during the EXPORTS campaign within an anticyclonic eddy, the trends that were observed here may in fact be a common feature of the region that is frequently overlooked or smoothed into long term averages but may have temporal carbon export implications.

Tracer experiments show that the four storms resulted in substantial advection of surface waters (ranging from 23-73% flushing, i.e., the amount of time required for a current driven by Ekman transport to displace a 15 km radius circle; Johnson et al., preprint) and changes in the mixed layer depth that ranged from 25.5 to 46.5 m. The most impactful of these storms was storm 1, occurring from JDs 127 – 130 and causing the most substantial advection and mixed layer depth change of the observational period (Johnson et al., preprint). Assuming this event stimulated our mid-cruise peaks in NPP and new production through enhanced nutrient availability, changes in the light field, or changes to community composition, we can estimate the impact of these episodic events by removing these enhanced production days (JD 132 for new production and JD 134 for NPP) and evaluating how much our integrated rates change. For NPP, elevated ^13^C-ρDIC on JD 134 increased integrated cruise NPP by 164 mmol C m^−2^ d^−1^, and for new production, elevated ^15^N-ρNO_3_^−^ on JD 132 increased the value by 2.89 x 10^4^ nmol N m^−2^ d^−1^, or 191 mmol C m^−2^ d^−1^. These values are equivalent to 8.4% and 13.2% of total cruise NPP and new production, respectively, consistent with results of previous studies which find that the contribution of episodic deep mixing events to total mixed layer production averages around 10%, but can be as high as 110% (Penta et al., 2021). Both 164 mmol C m^−2^ d^−1^ and 191 mmol C m^−2^ d^−1^ exceed maximum daily integrated values for both NPP and new production. The important role of mixed layer dynamics in primary production and biomass accumulation has been frequently reported (Dall’Olmo et al., 2016; Penta et al., 2021), but this study presents an added dimension: substantial changes to the mixed layer could play an important role in temporarily sustaining bloom-associated primary production, preventing a completely monotonic bloom dissipation.

Despite the roles of advection and mixed layer fluctuations, we continue to sample one water mass throughout the majority of the observation period, and underlying these physical dynamics, the typical phytoplankton taxonomic transition within the eddy following a diatom bloom appears to occur and coincides with changes in the pigment-based size class distribution (Fig 4; Henson et al., 2012b). The HPLC results suggest a dominant proportion of the community (on average 89%) was composed of members of the red algal lineage during the observational period with noteworthy fluctuations in pigments associated with diatoms (Fuco) and haptophytes (HexFuco; Figure 4). These transitions are consistent with observations collected from onboard imaging, which show a high abundance of diatoms in the beginning of the cruise before a transition to haptophytes and dinoflagellates (Perid, the marker pigment for some dinoflagellates, increased >5% over the observational period; H. Sosik, C. Roesler, and L. Karp-Boss, personal communication). These taxonomic trends are consistent with a shift away from a large-celled autotrophic community with high rates of primary production but surprisingly coincide with an increase in new production and NPP on JDs 132 and 134. The weighted Diagnostic Pigment Analysis shows that the temporal decline in the proportion of the large size fraction relative to the total NPP, new production, and Chl *a* concentrations is substantially higher (37%, 42%, and 37%, respectively) than the decrease in the relative proportion of microphytoplankton over time (16%). This result may suggest that the large cells still make up a substantial proportion of the community, but they are not actively growing or taking up as much nutrients in the latter half of the observational period relative to the beginning. This speculation is supported by the uncoupling of ^32^Si-ρSi (which tends to be associated with larger diatoms) relative to ^13^C-ρDIC (which encompasses all phytoplankton size fractions; Fig 10A). The uncoupling of ^32^Si-ρSi and ^13^C-ρDIC within the large size-fraction would suggest cells are enhancing silicification at a faster relative rate than NPP, making cells denser and heavier and thus, more likely to sink out of the euphotic zone, enhancing export. This notion is consistent with sediment trap and optical observations that suggest concentration of large particles are being exported from the upper water column to 500 m depth increases over the observation period (D. Siegel, personal communication).

Our high cruise average f-ratios suggest that nitrate is the dominant form of nitrogen used in primary productivity within the system consistent with nitrate concentrations remaining between 4 and 6 µmol L^−1^ during the cruise. Interestingly, while all size fractions exhibit f-ratios >0.5, the small size-fraction had a slightly higher average f-ratio (+0.04) than the larger size fraction. This finding is contradictory to the commonly observed trend that new production tends to be dominated in phytoplankton taxa typically >5 µm that are most commonly diatoms. Additionally, we observed substantially higher biomass in the large size fraction. This result likewise supports the idea that despite sustained biomass, the larger cells engaged in less production over the observational period. Evaluating the temporal trends shown on Figure 8, we see that the higher f-ratio in the small size-fraction is likely driven by variable rates of new production and consistently low rates of regenerated production relative to the comparatively stables rates of new production in the large size fraction. Given the scarcity of in situ data on regenerated production and f-ratios from this, it is difficult to evaluate whether our observations in this study are common or whether this trend is driven by the phytoplankton taxonomic assemblage shifts (Fig. 4) and low nutrient concentrations, or a combination of both. Despite being in non-steady state conditions, our f-ratios are consistent with Thorium-210 isotope data which suggest substantial export of carbon (>300 mmol C m^−2^) from the surface ocean over the observational period (S. Clevenger and K. Buesseler, personal communication).

While carbon and nitrogen uptake dynamics are uncoupled over various portions of the observation period, there was not a consistent pattern of elevated PC:PN ratios as previously observed in this region (Fig. S4; Fig. S5). The moderate to frequently lower than Redfield ratio stoichiometry suggests that carbon overconsumption was not occurring in the autotrophic community during our sampling. While there could be a number of potential explanations for this result, one possible reason is that ambient nitrate concentrations remained relatively high, unlikely reaching limiting concentrations for the majority of the phytoplankton assemblage. As diatoms made up a majority of the community when production and biomass were highest, silicic acid was more likely to be the primary limiting nutrient, rather than nitrogen, and silicic acid concentrations were routinely measured to be <2 umol L^−1^ in the euphotic zone. Silica limitation is supported by the inaccuracy of the traditional 0.13 Si to C conversion factor that often drove the percent siliceous NPP greater than 100%. Severe silica limitation in diatoms has been shown to exhibit a differential response to nitrate limitation (Brzezinski and Nelson, 1996; De La Rocha and Passow, 2004). Additionally, lowered C:N ratios in the North Atlantic have been reported as nitrate limitation subsided following deep mixing events (Penta et al., 2021).

Further evidence exists to suggest nutrient limitation may be a significant control on primary production, particularly within the small size-fraction. Extremely elevated PC:Chl *a* ratios were observed on JD 126-128. Elevated PC:Chl *a* ratios in phytoplankton have frequently been reported in systems experiencing severe nutrient, primarily iron, limitation or co-limitation (Sunda and Huntsman, 1997). Extremely low silicic acid concentrations may contribute to these elevated ratios; particularly, given the discontinuity between average to below-average PC:PN ratios over the course of the observation period. However in contrast to what we observed at the beginning of the sampling period, elevated PC:Chl *a* ratios were also observed near the end of sampling in both size fractions and during a period when overall NPP was low. These high ratios coincide with a time when a substantial proportion of total POC appears to be detrital (J. Graff, personal communication). Additionally, during this period, substantial POC was shown to be sinking out from the surface ocean to depths >400 m (D. Siegel, personal communication). Taken together, these results suggest differing factors causing extremely elevated PC:Chl *a* ratios at the beginning and end of the observation period, each of which has important ecological and biogeochemical consequences.

Additionally, we see evidence to suggest that storm events caused deepening of the mixed layer which upwelled waters containing higher nutrient concentrations into the euphotic zone. This result is particularly evident in our Z-scored data on JD 132 (Figure 11). From sample days 128 and 132, the mixed layer deepened by approximately 27 m, Si(OH)_4_ concentrations increased by 1.05 µmol L^−1^, and NO_3_ ^−^ concentrations increased by 0.23 µmol L^−1^. Presumably, this event led to enhanced new production in the small cells (+63 mmol C m^−2^ d^−1^) and total new production (+72 mmol C m^−2^ d^−1^) as well as ^15^N-*V*NO_3_ ^−^. Here, we see slight decoupling between ^15^N-ρNO ^−^ and ^13^C-ρDIC since ^13^C-*V*DIC and NPP do not increase until the following sample day on JD 134. Previous studies have likewise reported a connection between turbulence-induced mixing and enhanced new production with noteworthy connections to carbon export (Dall’Olmo et al., 2016; Penta et al., 2021). Additionally, the physical nature of eddies and their associated water movement have been shown to provide enhanced macronutrients, underscoring the potential influence lateral transport may have played in these dynamics (Garside and Garside, 1992; Hartman et al., 2010).

### 4.3 Global comparison and context

The two-endmember approach of the EXPORTS program and array of parameters collected via near identical methods in the two ocean basins allows for a nearly direct comparison of the various controls on primary production within these vastly different systems. Unsurprisingly, the drastically different ecological, physical, and biogeochemical attributes of the two basins led to differing key characteristics and drivers between the systems as shown through the different branching and clustering patterns of parameters in the dendrograms (Fig. 12). At the North Pacific EXPORTS site, parameters were less separated by size class relative to the North Atlantic (Fig. 12A). Figure 12A divides into two primary branches with the majority of biomass (Chl *a*, PC, and PN), NPP, new production, f-ratios, and GPP : NPP along with T, PAR, and MLD all on one branch. All four macronutrients and salinity link together on the second primary branch with *V*DIC and *V*NO_3_^−^ of both size fractions closely related (linkage distance = 13.5). This would suggest a tighter relationship between macronutrients and assimilation rates than phytoplankton standing stock or production itself. In Fig. 12B, all parameters related to the uptake of silicic acid (although not Si(OH)_4_ concentration itself) branch out on the second primary branch whereas macronutrients, salinity, and small size-fraction ρDIC : ρNO_3_ are on the first primary branch. The linkage distance between NO_3_^−^ concentration and small size-fraction ρDIC : ρNO_3_^−^ is the smallest of any two parameters at 1.20. This is consistent with previous classifications of the region which characterize the region as being dominated by small cells engaging in the majority of production and high and relatively stable macronutrient concentrations (Harrison 2002; Meyer et al., 2022).

**Figure 12.**
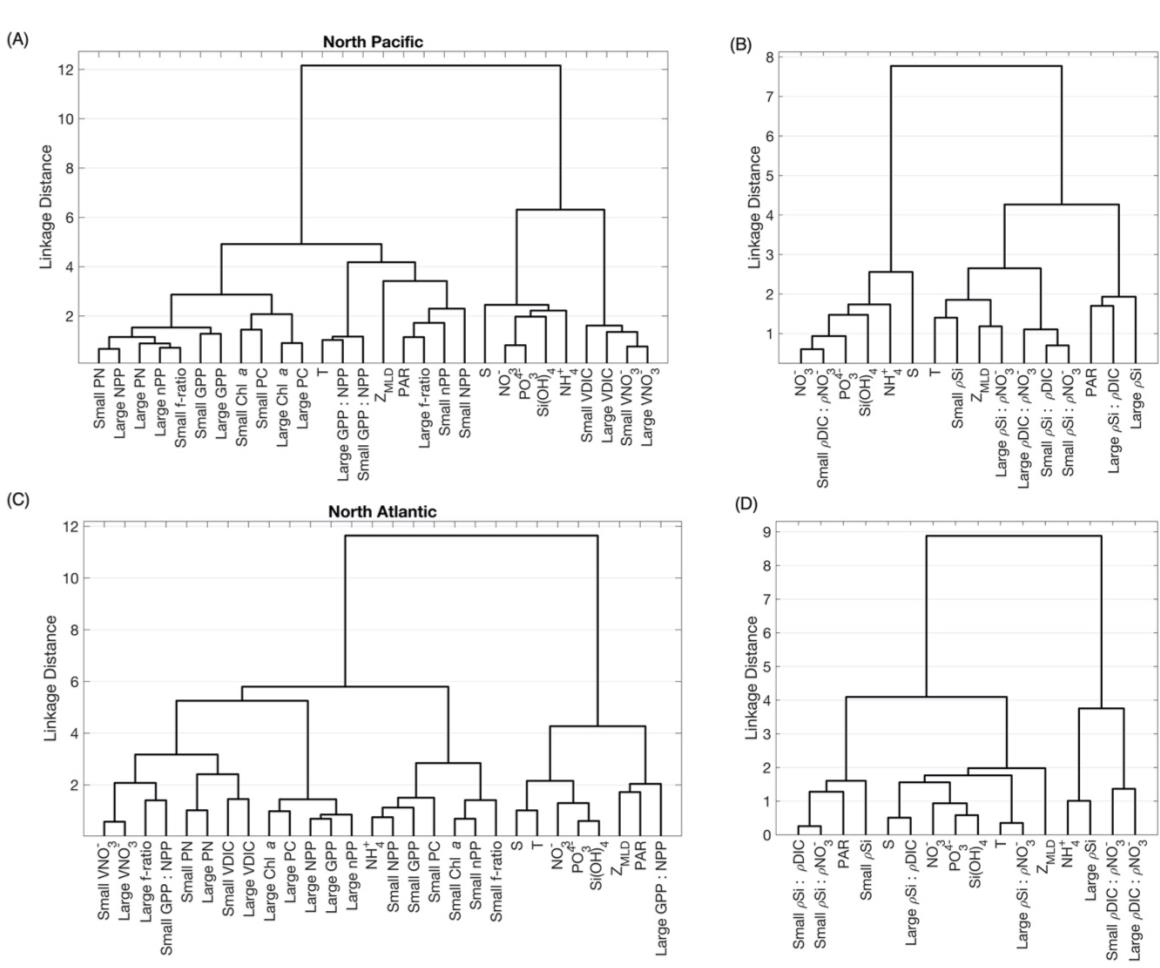
Dendrograms of environmental parameters, phytoplankton parameters, and production for samples collected in the North Pacific and North Atlantic. Dendrograms (Euclidean distance linkages) of environmental and phytoplankton biomass, production, and assimilation parameters from the EXPORTS (A) North Pacific and (C) North Atlantic campaigns. Environmental parameters, ρSi, and physiologically informative ratios were separated into (B) for the North Pacific and (D) for the North Atlantic due to lower data resolution for ρSi which could confound linkage distance of variables that were sampled at higher resolution.

At the North Atlantic EXPORTS site, we see a greater separation of environmental parameters into the second primary branch (Fig. 12C). Noteworthy exceptions are NH_4_^+^ which links within a branch entirely comprised of small size-fraction biomass and production parameters as well as large size-fraction GPP : NPP which links within the environmental parameters branch, most closely with MLD and PAR. The close linkage (linkage distance = 3.4) between MLD and PAR may reflect enhanced cloud cover and mixed layer variability driven by the storm events. In the North Atlantic, we see greater separation by size class particularly in Chl *a*, PC, and the production rates suggesting greater size dichotomy than was evident in the North Pacific. The exceptions to this are PN, *V*DIC, *V*NO_3_^−^, the large size fraction f-ratio, and the small GPP : NPP which link together in a subbranch. In the ratios dendrogram (Fig. 12D), we see size appears to be a driving influence separating like ratios, ie., small size-fraction ρSi : ρDIC is more closely linked to small size-fraction ρSi : ρNO_3_ and large size-fraction ρSi : ρDIC is more closely linked to large size-fraction ρSi : ρNO_3_^−^ (Fig. 12D). Interestingly though, small and large size-fraction ρDIC : ρNO_3_^−^ branch out independently on the second primary branch, thus suggesting they are more closely related to NH ^+^ and large size-fraction ρSi. This may suggest a separation of the dynamics of siliceous phytoplankton, which appear more coupled to PAR, salinity, temperature, and most macronutrient concentrations, from the remaining non-siliceous phytoplankton community. However, it is curious that large size-fraction ρSi separates out onto the second branch. This may support the notion of the uncoupling of ρSi from ρDIC and ρNO ^−^ dynamics as diatoms become more heavily silicified without engaging in substantially more NPP.

Size-fractions and environmental parameters play key, yet differing, roles in both systems. The North Pacific represented a physically stable system where slight changes in nutrient dynamics impacted phytoplankton assimilation which translated into modest changes in production but not necessarily biomass (Fig. 12A; Meyer et al., 2022). Whereas in the North Atlantic, the degree of change in mixed layer depths and nutrient concentrations was larger, leading to greater changes in production and phytoplankton standing stock. The dendrograms here highlight these environmentally important relationships while masking the degree of change occurring per system. This type of analysis can help establish linkages and drivers but their net impact of sed drivers need to be evaluated independently to assess their influence on large scale ecosystem dynamics and biogeochemistry.

### 4.4 The importance of size on the global scale

Size is well understood as a master trait in ocean ecology as well as a key variable in oceanic modelling (Blanchard et al., 2017). Results from the North Pacific and North Atlantic EXPORTS campaigns show that the role of cell size in biogeochemical cycling and export processes is complex. In the North Pacific, the community at Ocean Station Papa was dominated by small cells that were responsible for the majority of NPP, which fed into an inefficient, regenerative food web with low carbon export (Meyer et al., 2022; McNair et al., preprint). However, periods of higher-than-average NPP were driven by enhanced new production in both small and large cells, reflecting an important role for the large size fraction, as well as the small size-fraction, in shaping ecosystem dynamics. Conversely, in the North Atlantic, the majority of biomass and NPP were in the large size-fraction. However, NPP, new production, and regenerated production did not exhibit the gradual decline but rather temporarily declined before exhibiting high production rates again on JD 132 and 134 (Fig 8). The sustained production rates appear disproportionally driven by new production occurring in the small cells (Fig 8C; Fig 11). Some of this may be due to mixing from the first storm event as well as enhanced grazing by microzooplankton that is observed mid-cruise (JDs 138 – 145; H. McNair, personal communication). Additionally, the divergent production response between large and small size fractions during this time is represented in the opposite trends of parameters such as *V*DIC, ρDIC : ρNO_3_, ρSi : ρDIC, and ρSi : ρNO_3_ during this time (Fig. 11). In both systems, an important pattern emerges within the size fractions: one size fraction dominates biomass and production dynamics, while the other size fraction plays an important role in production dynamics, but not biomass, during periods when conditions (i.e., nutrient concentrations, predation, ballasting, etc.) temporarily become favorable for this size fraction.

This pattern is more likely to be upheld in systems that fall on the extreme ends of the primary production spectrum. These systems are known to maintain communities that are predominantly composed of one size fraction over the majority or over specific periods of the year (Harrison, 2002; Marchetti et al., 2006; Henson et al., 2012; Nowicki et al., 2022). Such dichotomy of size fractions in systems that fall within the middle of the production spectrum is less likely. However, these extreme systems are of disproportional importance when trying to understand drivers of carbon export variability and the implications under future climate change scenarios (Henson et al., 2021; Benedetti et al., 2021). As such, representing these relationships accurately in climate models is critical. Some advanced models are able to account for size class differences (Archibald et al., 2019; Henson et al., 2021; Nowicki et al., 2022). However, these models are computationally difficult and maintain high uncertainties. Until we can better resolve the drivers and relative importance of size-fractionated primary production in situ, the accuracy and applicability of these models will remain constrained.

## 5. Conclusions

Our study highlights the impact substantial changes in mixed layer depth can have on the primary production dynamics during the decline of the annual North Atlantic spring phytoplankton bloom and the potential impacts for carbon export in this system. Our data support the findings of other studies from the North Atlantic (Hartman et al., 2010) and further characterize this region as having high f-ratios and high rates of primary production occurring in the large size fraction at the beginning of the observational period before a transition to near equal rates of production in the small and large size fractions following substantial mixing events. This study highlights the importance of cell size and macronutrient availability in driving net trends. Silicic acid concentrations appear to be a key control on siliceous phytoplankton biomass production with concentrations in the euphotic zone varying in relation to both uptake and mixed layer variability. Our results support further investigation into the role of size and the net impact of mixed layer variability (i.e., the mixed layer pump) in this region and the need for a better understanding of the role of eddies, storm events, and advection on primary production and export trends during the North Atlantic bloom and how these impacts can be incorporated into broad scale biogeochemical models.

## Data Accessibility Statement

All data presented here are available at the SeaWiFS Bio-optical Archive and Storage System (SEABASS; seabass.gsfc.nasa.gov/cruise/EXPORTSNA).

## Acknowledgements

We would like to thank the EXPORTS project leaders, D. Siegel and I. Cetinic, the chief scientists, D. Steinberg, J. Graff, D. Siegel, and C. Lee, and the captains and crews of the RRS *James Cook* and RRS *Discovery*, and the entire EXPORTS team. In particular, we would like to thank H. McNair, D. Fontaine, and S. Traylor for assistance in the field and the EXPORTS Hydro team for nutrient data.

## Funding

This work was supported by NASA Grant 80NSSC17K0552 to AM, SG, and NC and NSF Grant OCE-1756442 to MB. SK was supported by NASA Grant 80NSSC17K0692 to D. Siegel, and NP was supported by NASA Grant 80NSSC18K143 to A. Santoro.

## Competing Interests

The authors declare no competing interests.

## Notes

https://seabass.gsfc.nasa.gov/

### Competing Interest Statement

The authors have declared no competing interest.

