## Supplemental Figures and Tables for "Primary production dynamics during the decline phase of the North Atlantic annual spring bloom"

Supplemental Material

**Supplemental Table 1.** Pigments used and their distribution across key phytoplankton functional groups. Pigment names are shortened from fucoxanthin (Fuco), peridinin (Perid), 19'-butanolyoxyfucoxanthin (ButFuco), 19'-hexanoyloxyfucoxanthin (HexFuco), alloxanthin (Allo), chlorophyll b (Chl b), and zeaxanthin (Zea). Table modified from Kramer and Siegel (2019).

| | Fuco | Perid | But-Fuco | Hex-Fuco | Allo | Chl b | Zea |
|---|---|---|---|---|---|---|---|
| Diatoms | | | | | | | |
| Dinoflagellates | | | | | | | |
| Crysophytes | | | | | | | |
| Pelagophytes | | | | | | | |
| Haptophytes | | | | | | | |
| Cryptophytes | | | | | | | |
| Prasinophytes | | | | | | | |
| Euglenoids | | | | | | | |
| Chlorophytes | | | | | | | |
| *Trichodesmium* | | | | | | | |
| *Synechococcus* | | | | | | | |
| *Prochlorococcus* | | | | | | | |

| | |
|---|---|
| | Unique |
| | Always present |
| | Often present |
| | Rarely present |
| | Trace |
| | Not present |

Supplemental Figures:

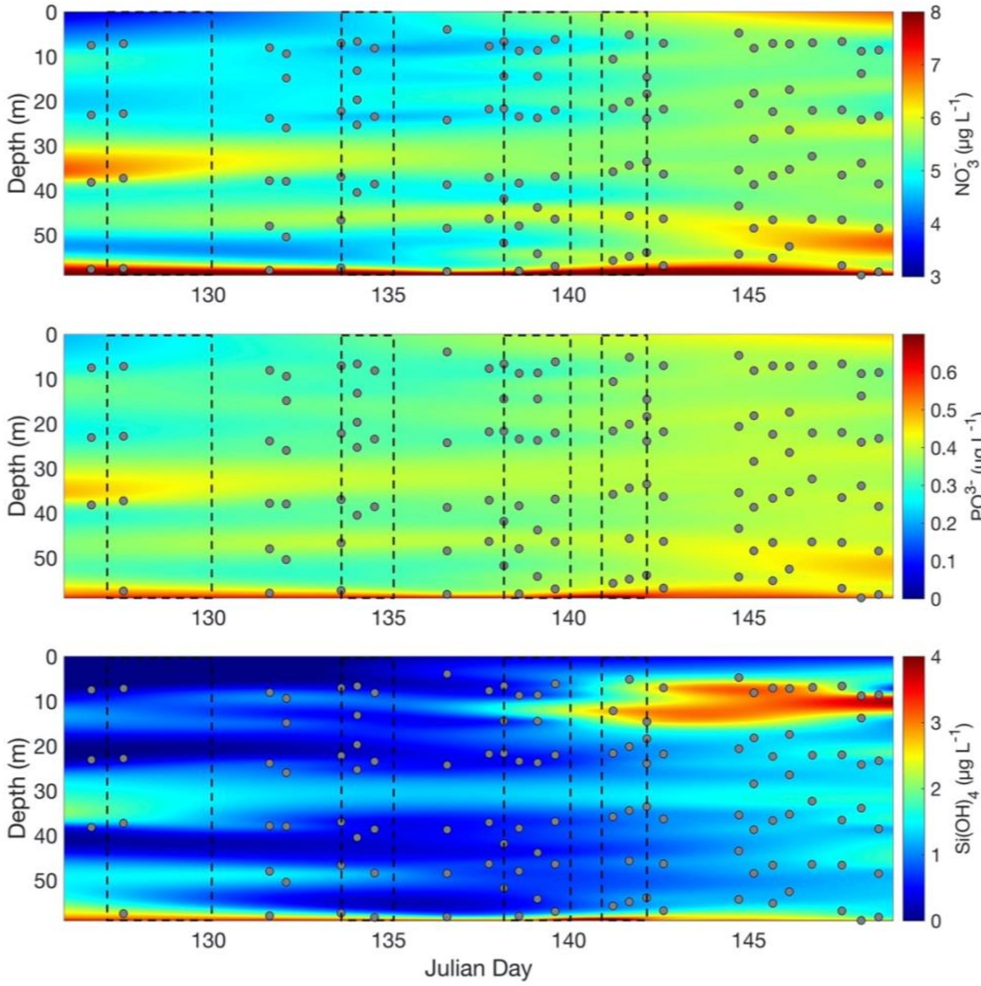


**Figure S1. Discrete macronutrient concentrations through time.** Discrete, euphotic zone nitrate ($NO_3^-$), phosphate ($PO_4^{3-}$), and silicate ($Si(OH_4)$) concentrations by Julian day. Units are in µg $L^{-1}$. Black boxes indicate storm events.

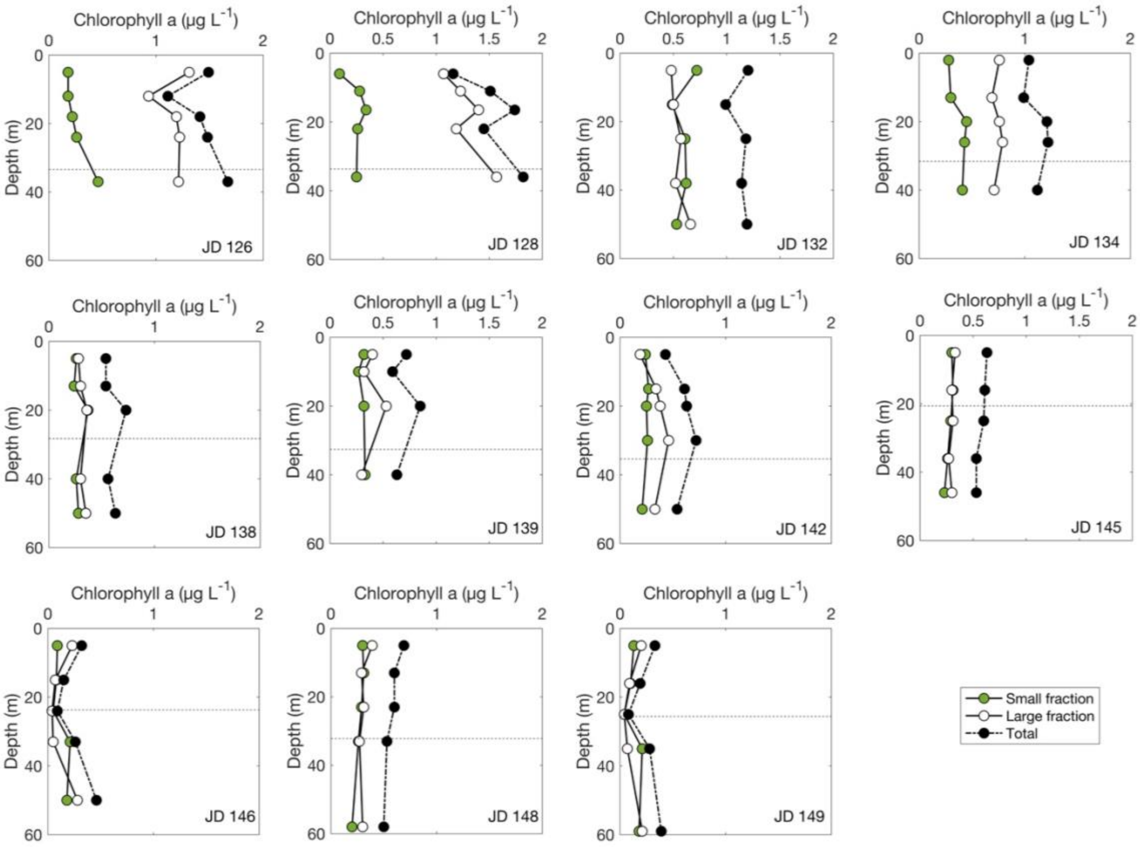


**Figure S2. Size-fractionated chlorophyll a concentrations**. Discrete size-fractionated chlorophyll *a* concentration (μg $L^{-1}$) by sample depth (m) at up to five depths throughout the euphotic zone. Small size-fraction (green circles) refers to cells between 0.7-5 μm and large-size fraction (white circles) refers to cells ≥5 μm. Totals (black circles) refer to cells >0.7 μm. Dashed lines indicate mixed layer depths. Each sample represents the means of triplicates. Standard deviations were small, typically 16.1% and 18.0% of the means for small and large size fractions, respectively.

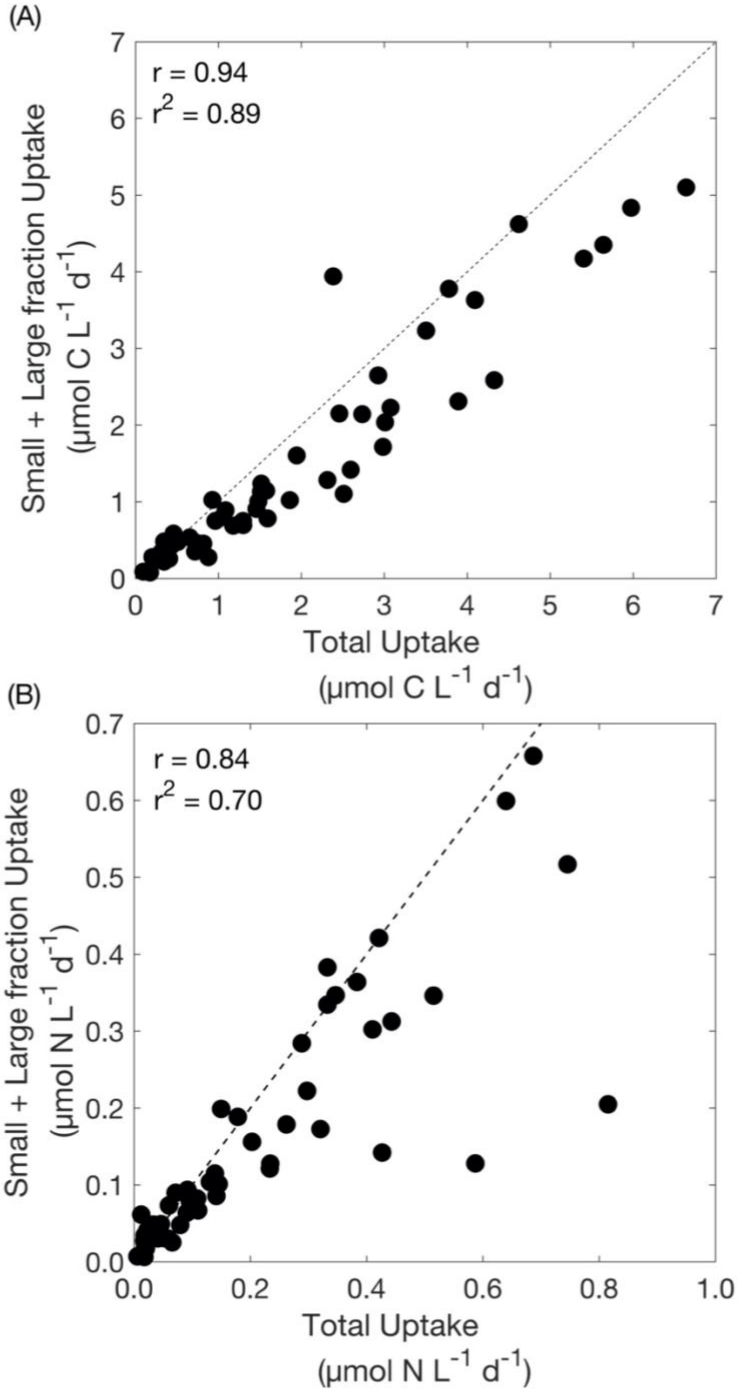


**Figure S3. Summed size fractionated versus total uptake rates.** Discrete summed (small + large) fraction carbon uptake (μmol C $L^{-1}$) and nitrate uptake (μmol N $L^{-1}$) versus total fraction (A) carbon and (B) nitrogen uptake, respectively. The small size fraction refers to particles <5 μm, and the large size fraction refers to particles ≥5 μm. The dashed line indicates a 1:1 line. R and $R^2$ values per regression are indicated.

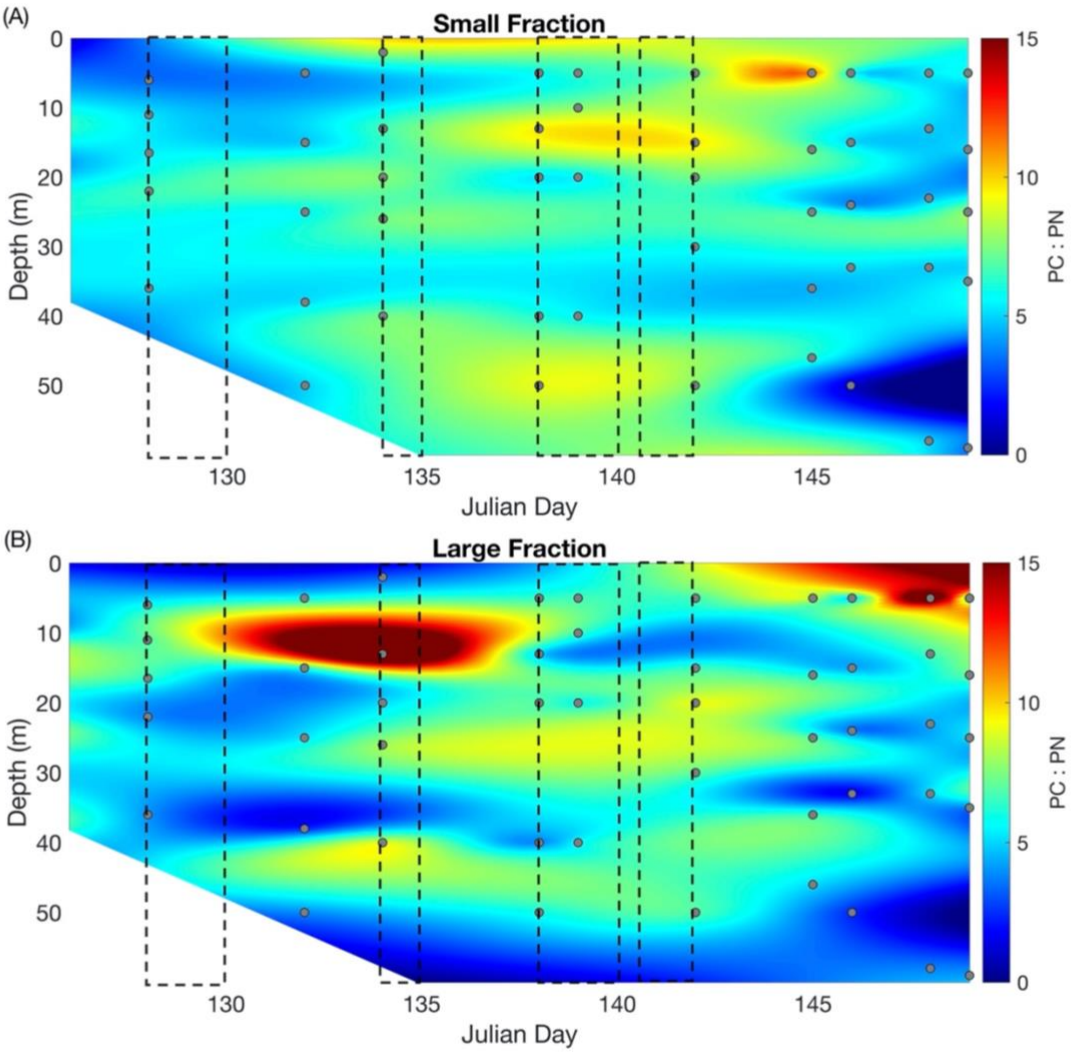


**Figure S4. Size fractionated PC:PN ratios.** Discrete size fractionated particulate carbon to particulate nitrogen (PC:PN) ratios for (A) small (<5 µm) and (B) large (≥5 µm) cells throughout the euphotic zone and by Julian day. Units are µmol C $L^{-1}$ : µmol N $L^{-1}$.White regions indicate areas without data collected. Black boxes indicate storm events. Ratios between samples are interpolated and thus introduce some uncertainty to those estimates.

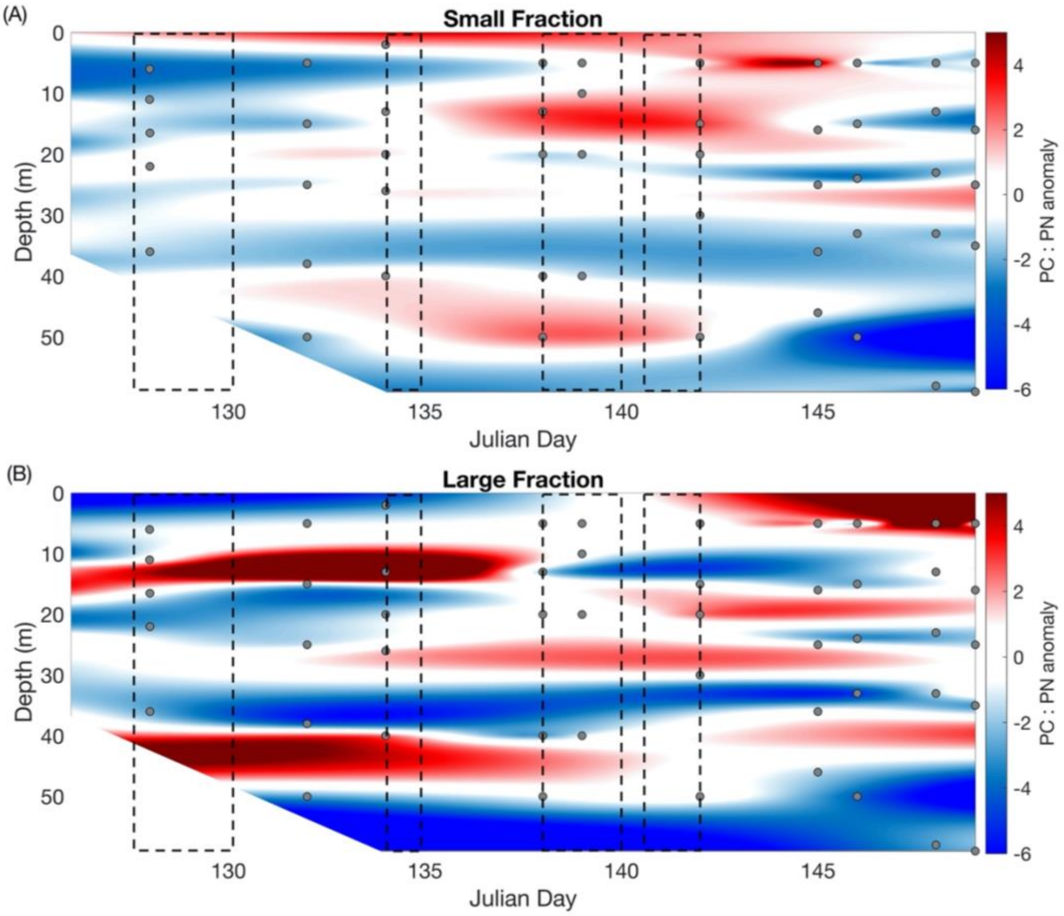


**Figure S5. Size fractionated PC:PN ratio anomalies.** Discrete size fractionated particulate carbon to particulate nitrogen (PC:PN) ratio anomalies for (A) small (<5 µm) and (B) large (≥5 µm) cells throughout the euphotic zone and by Julian day. Anomalies are calculated as measured ratio using the canonical Redfield ratio (6.6C : 1N). Units are µmol C $L^{-1}$ : µmol N $L^{-1}$.White regions indicate areas without data collected. Black boxes indicate storm events. Ratios between samples are interpolated and thus introduce some uncertainty to those estimates.

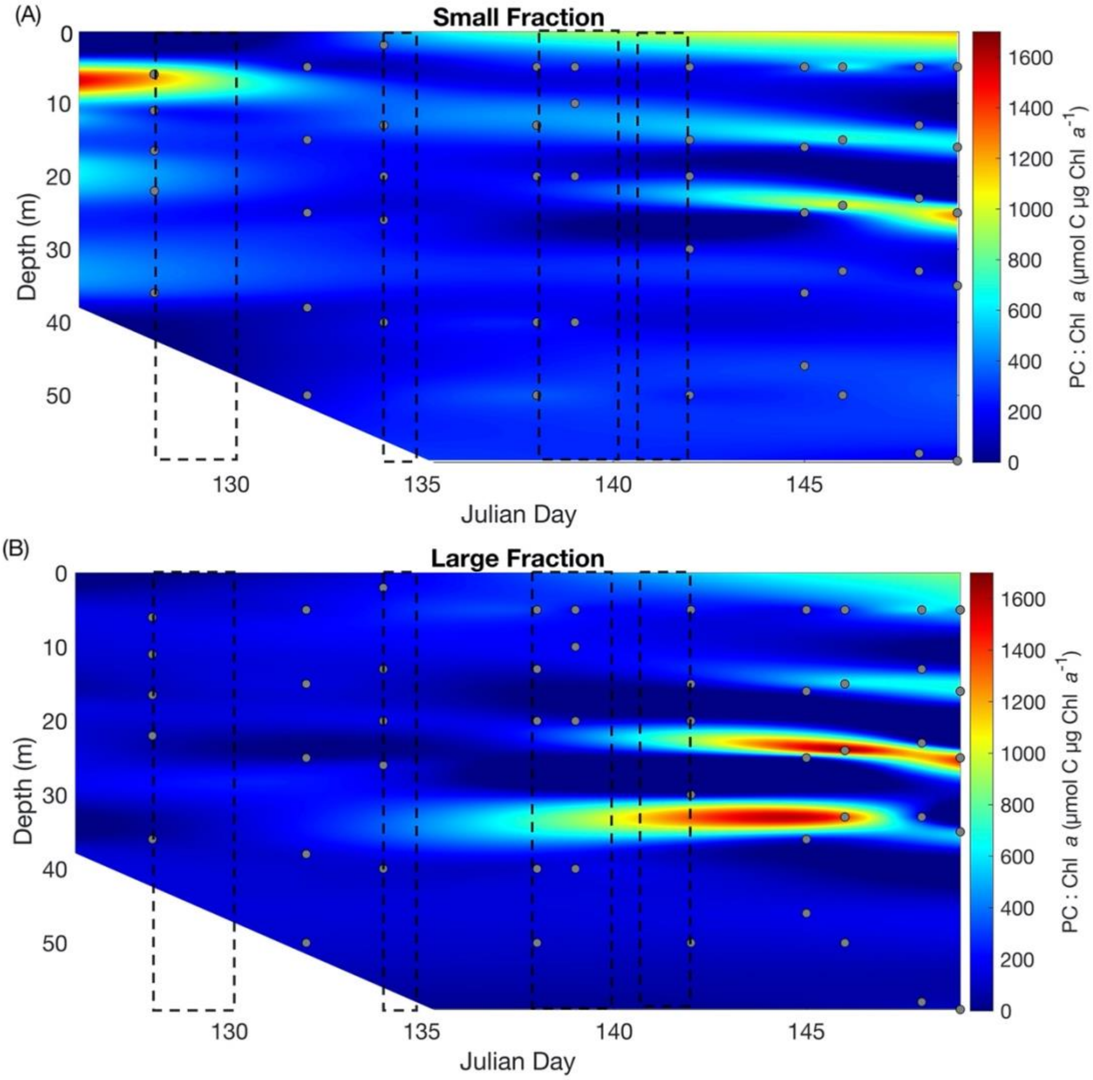


**Figure S6. Size fractionated PC:Chl *a* ratios.** Discrete size fractionated particulate carbon to chlorophyll *a* (PC:Chl *a*; µmol C µg Chl $a^{-1}$) ratio anomalies for (A) small (<5 µm) and (B) large (≥5 µm) cells throughout the euphotic zone and by Julian day. White regions indicate areas without data collected. Black boxes indicate storm events. Ratios between samples are interpolated and thus introduce some uncertainty to those estimates.

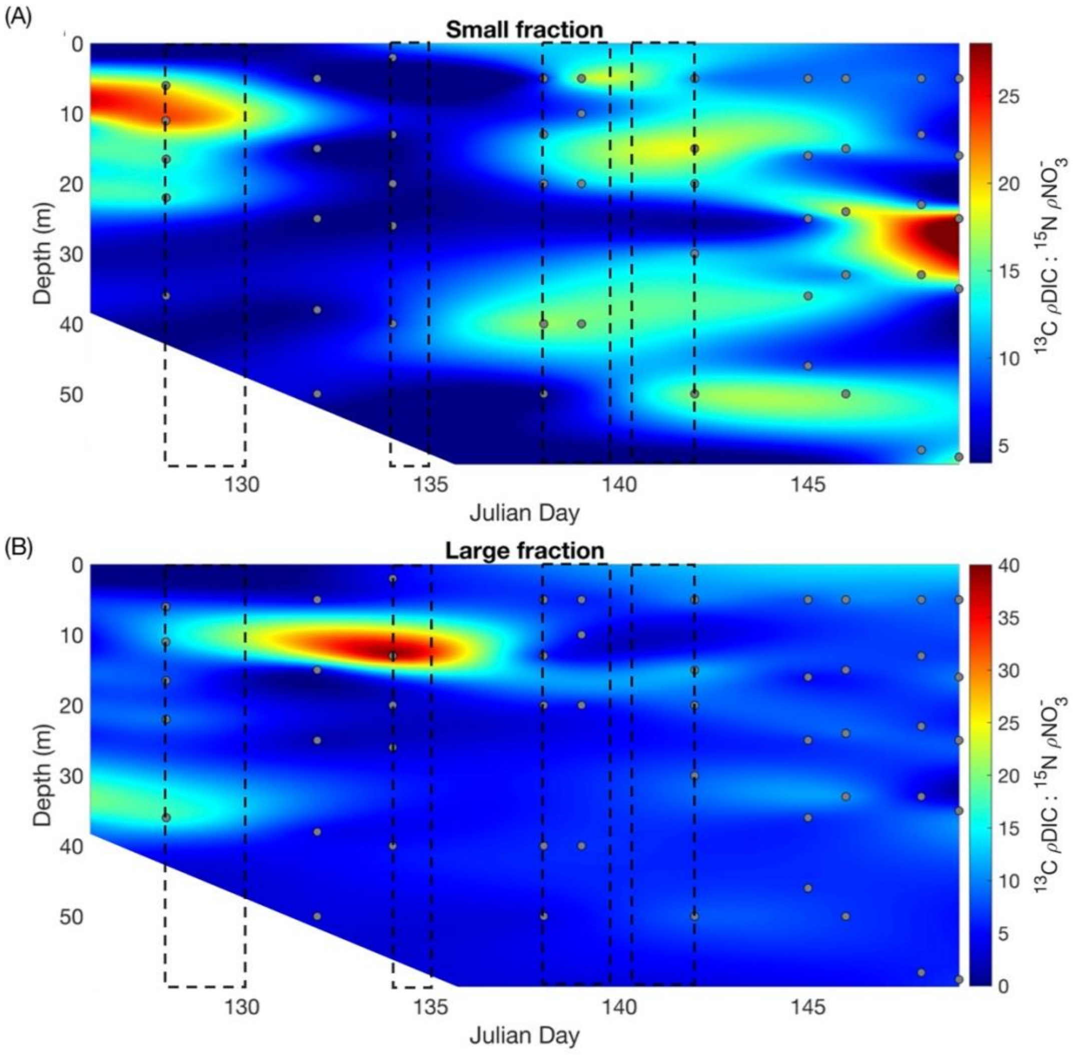


**Figure S7. Size fractionated carbon uptake to nitrate uptake ratios.** Discrete size fractionated carbon-13 uptake to nitrogen-15 uptake ($^{13}$C-ρDIC: $^{15}$N-ρ$NO_3^-$) ratios for (A) small (<5 µm) and (B) large (≥5 µm) cells throughout the euphotic zone and by Julian day. $^{15}$N-ρ$NO_3^-$ was converted to carbon units via the Redfield ratio (6.6C : 1N). White regions indicate areas without data collected. Black boxes indicate storm events. Ratios between samples are interpolated and thus introduce some uncertainty to those estimates.
